# Ex vivo stem-like cell families model evolution of glioblastoma therapeutic resistance

**DOI:** 10.64898/2026.04.02.716158

**Authors:** Marta Prelli, Francesca De Bacco, Elena Casanova, Salvatore Maniscalco, Gaia Biagioni, Gigliola Reato, Salma Mahmoudi, Raffaele Calogero, Mara Panero, Erika Boasso, Laura Casorzo, Giovanni Crisafulli, Alice Bartolini, Marco Macagno, Zachary D. Nagel, Luca Bertero, Paola Cassoni, Pietro Zeppa, Fabio Cofano, Diego Garbossa, Francesca Orzan, Carla Boccaccio

## Abstract

Glioblastoma (GBM) arises from stem-like cells (GSCs) that exhibit intrinsic therapeutic resistance and can be positively selected by treatment, rendering recurrent GBM intractable. Mechanistic dissection of therapeutic resistance evolution has been limited by scarce matched primary/recurrent tractable models. To address this gap, we developed “resistant GSC families”, a within-patient matched platform that models therapy-driven selection by ex vivo deriving, from the same primary whole GBM, temozolomide (TMZ)- and ionizing radiation (IR)-selected GSCs, alongside a treatment-naïve control (CTRL-GSC). This design enables pressure-specific dissection of resistance evolution, separating chemotherapy- and radiotherapy-associated genetic alterations and adaptations that are often confounded in primary-versus-recurrent comparisons. Using this framework, we link TMZ-driven relapse-like states to either mismatch repair (MMR)-dependent stable resistance or O6-methylguanine-DNA methyltransferase (MGMT)-independent drug tolerance, and identify adaptive DNA damage response and cell-cycle changes as a route to increased radioresistance. Across pressures, treatment-emergent GSCs accumulate chromosomal alterations and exhibit adaptive phenotypic remodeling, including increased receptor tyrosine kinase activity. Resistant GSC families represent a model enabling mechanistic studies and hypothesis-driven testing of strategies aimed at preventing or treating GBM recurrence.

## INTRODUCTION

Glioblastoma (GBM) is the most aggressive primary brain tumor in adults, characterized by diffuse infiltration, rapid progression, and near-universal recurrence despite maximal treatment (Weller *et al*, 2024; Wen *et al*, 2025). Current standard-of-care therapies, including surgical resection followed by radiotherapy and temozolomide (TMZ) chemotherapy, offer only modest survival benefits, with median overall survival rarely exceeding 15 months (Weller *et al*., 2024; Wen *et al*., 2025). A major barrier to durable responses is the intrinsic and acquired resistance of a subpopulation of tumor cells with stem-like properties—glioblastoma stem-like cells (GSCs)—which contribute to tumor maintenance, therapy resistance, and relapse (Bao *et al*, 2006; Chen *et al*, 2012; Gimple *et al*, 2022; Osuka & Van Meir, 2017; Sloan *et al*, 2024).

Previous studies, including our own, have shown that distinct GSC subpopulations coexist within the same tumor and display marked heterogeneity, including potential differential sensitivity to cytotoxic agents and ionizing radiation (IR) (De Bacco *et al*, 2023b; Meyer *et al*, 2015; Piccirillo *et al*, 2015; Piccirillo *et al*, 2009; Suvà & Tirosh, 2020). This intra-tumoral heterogeneity provides a substrate for therapeutic selection, whereby GSC subclones with enhanced resistance may be positively selected under treatment pressure, ultimately driving recurrence (Nicholson & Fine, 2021). However, the mechanisms by which therapeutic pressure shapes the evolution of GSC resistance traits remain poorly understood, largely because matched primary and recurrent samples are rarely available, as relapsing GBMs are seldom resected, limiting generation of robust, tractable models that recapitulate resistance development (Lucchini *et al*, 2025).

To address these gaps, we developed an *ex vivo* protocol that enables parallel isolation of therapy-naïve GSCs (CTRL-GSCs) and their treatment-selected counterparts—TMZ-GSCs and IR-GSCs—directly from primary GBMs recovered and processed to maximally preserve heterogeneity of the initial whole cell population. We used these “resistant GSC families” (res-GSC families) to dissect genetic, phenotypic, and functional changes associated with TMZ or IR therapeutic pressure and the development of pressure-specific resistance. GSCs emerging from ex-vivo therapies recapitulated established mechanisms of acquired TMZ resistance, including mismatch-repair gene inactivation, supporting the relevance of the model to patient disease. In addition, res-GSC families revealed still poorly understood resistance features, including persister-like states, incremental radioresistance linked to increased DNA damage response efficiency, and cross-treatment increases in karyotypic alterations and receptor tyrosine kinase (RTK) activity. This ex vivo model enables within-patient matched, experimentally tractable studies and supports controlled mechanistic analyses to identify vulnerabilities that may inform more effective therapeutic strategies at the primary or recurrent stage.

## RESULTS

### Ex vivo derivation of res-GSC families models therapeutic selection and evolution from primary to recurrent GBM

We developed a protocol to *ex vivo* select GSCs that model those emerging after *in vivo* therapy and potentially driving tumor recurrence (Fig. 1A). To this end, we collected near-complete GBM cell populations surgically removed using an aspirator, processed them into single-cell suspensions, and cultured them in an enriched GSC medium (EFPH20) containing EGF, FGF2, PDGFBB and HGF (each at 20 ng/mL), based on a protocol we previously established to investigate GSC heterogeneity in individual GBMs (De Bacco *et al*, 2023a; De Bacco *et al*., 2023b). The growth factors in EFPH20 reflect those abundant in the brain and GBM microenvironment (De Bacco *et al*, 2023a; Sjöstedt *et al*, 2020), support key survival pathways involved in therapy resistance (Gimple *et al*., 2022), and are well suited to preserve the original GSC heterogeneity before therapeutic selection. After a brief recovery period (∼20 days) to remove cell debris, single-cell cultures derived from each GBM were exposed in parallel to selective therapeutic pressures: temozolomide (TMZ) at 50 μM every three days for three cycles, or ionizing radiations (IR) at 2 Gy per day for five consecutive days. A third culture was left untreated to serve as the control, representing treatment-naïve GSCs residing in primary GBM (Fig. 1A).

**Fig. 1.**
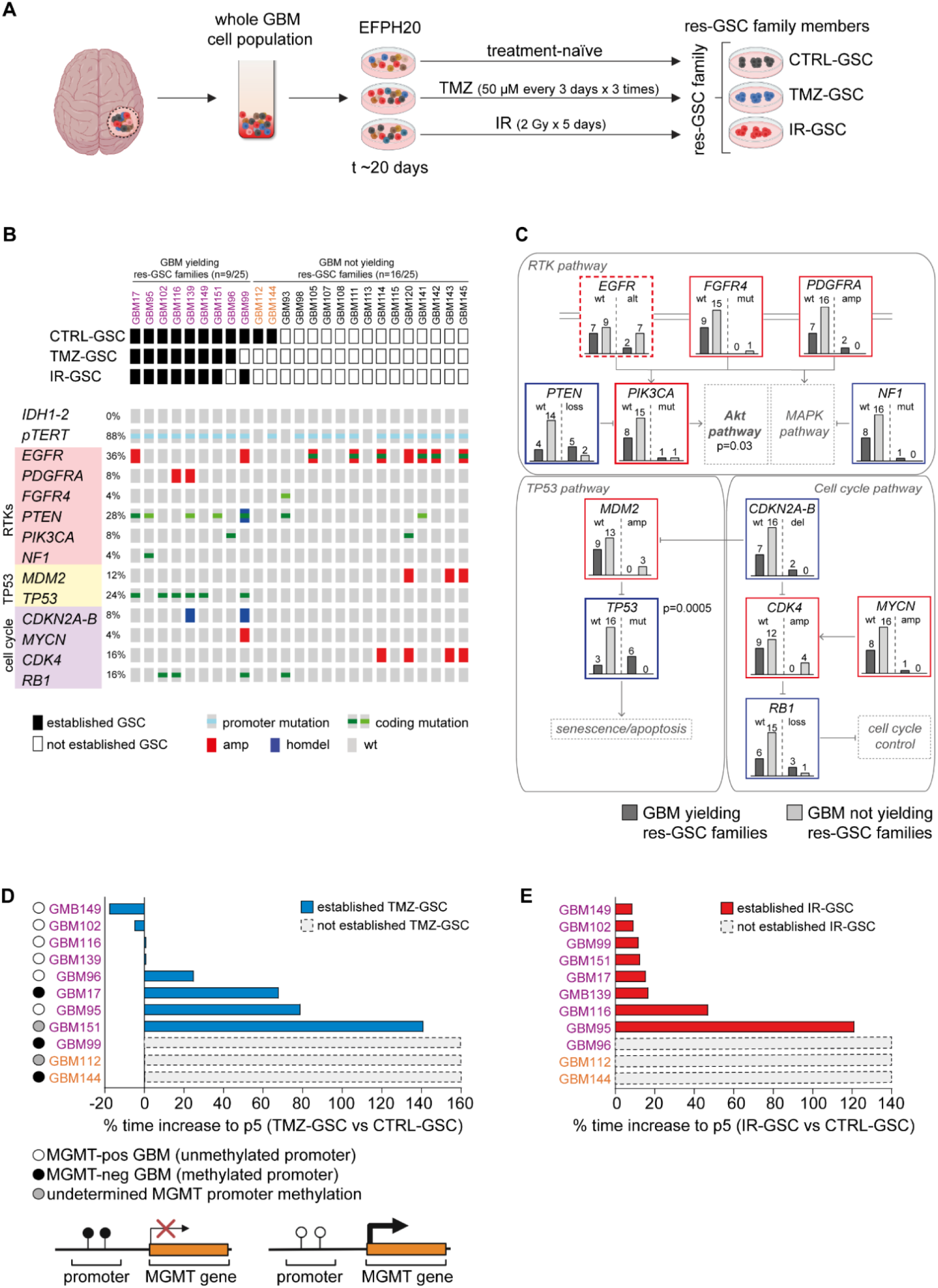
*Ex vivo* derivation of res-GSC families models therapeutic selection and evolution from primary to recurrent GBM. (**A**) Schematic of ex vivo treatment of whole GBM cell populations and generation of res-GSC families with chemo- or radiotherapy. Whole GBM cell suspensions are cultured in stem cell medium (EFPH20) supplemented with EGF, FGF2, PDGFBB, and HGF (20 ng/mL each) and exposed in parallel to temozolomide (TMZ) or ionizing radiation (IR) as indicated. Each res-GSC family includes TMZ-GSC, IR-GSC, and an untreated CTRL-GSC member. Families were considered established after passage 10. (**B**) Genetic profiling of original GBMs (n = 25) grouped by ability to generate res-GSC families under therapeutic pressure (columns). Top rows: res-GSC family members derived from each GBM (black squares). Bottom rows: altered genes and frequencies in original GBMs assessed by the NGS GBM-target panel. p*TERT* mutation was evaluated by ddPCR. Purple: GBMs yielding res-GSC families; light orange: GBMs yielding only CTRL-GSCs. (**C**) Associations between genetic alterations in original GBMs and res-GSC family derivation. Bars indicate the number of GBMs that are wild-type (wt) or altered (alt) for each gene and that did (dark gray) or did not (light grey) yield res-GSC families. Thicker dotted outline: tendency to anti-correlation between *EGFR* alterations and res-GSC family generation (p=0.40). *PTEN* loss and *PIK3CA* alterations showed gene-level trends (p = 0.06 and p > 0.99, respectively), while combined AKT-pathway alterations (*PTEN* and *PIK3CA*) were significantly associated with res-GSC family derivation (p = 0.03). *TP53* mutations were also significantly associated (p = 0.0005) (Fisher’s exact test). Red: oncogenes. Blue: tumor suppressor genes. (**D-E**) Derivation times of TMZ-GSCs (D, upper) and IR-GSCs (E) from GBM processing (as in **B**) to p5. Bars show the percentage increase in derivation time relative to the matched CTRL-GSC. Grey dotted bars: not derived GSCs. Absolute derivation times (days) are reported in Fig. EV2A. In (**D**), dots indicate the *MGMT* status of the original GBM (or of CTRL-GSC when the GBM status was unavailable). Lower panel: schematic of the functional relationship between *MGMT* promoter methylation and *MGMT* transcription (arrow).

The treatments protocols applied are clinically relevant, because they (i) mimic drug concentration measured in patients’ blood plasma and schedules used in patients (Ostermann *et al*, 2004; Stupp *et al*, 2005), and (ii) effectively distinguish primary sensitive from primary resistant GSCs, as previously assessed with irradiation experiments (De Bacco *et al*, 2016) and observed in TMZ dose-response *in vitro* experiments with an independent panel of GSC cultures (Fig. EV1A-C) (De Bacco *et al*., 2023b). Cells that survived these treatments - as well as treatment-naïve controls - gave rise to long-term propagating cultures that predominantly grew in suspension as neurospheres, although some formed adherent foci or exhibited intermediate growth patterns, and were considered as established at passage 10 (p10). From each GBM, we defined the group of established GSCs - including at least one therapy-selected culture - as a “res-GSC family”, with each individual culture referred to as a res-GSC family “member”. Specifically, these were designated as CTRL-GSC (treatment-naïve), TMZ-GSC (emerging after TMZ treatment), or IR-GSC (after IR treatment) (Fig. 1A). *In vitro* limiting dilution assays (LDA) showed a consistently significant increase in stem-like cells in IR-GSCs compared to CTRL-GSCs across all families, whereas TMZ-GSCs exhibited variable fractions of stem-like cells (Fig. EV1D-E).

The *ex vivo* derivation protocol was applied to 25 GBMs, i.e., grade 4 gliomas with IDH1/2 wild-type genes, as defined by histopathological diagnosis according to WHO 2021 CNS tumor classification (Table EV1) (Louis *et al*, 2021). Nine res-GSC families were successfully derived, seven of which comprising all three members, and two including only CTRL- and TMZ- or IR-GSCs (Fig. 1B; Table EV1). Among the remaining 16 GBMs, two cases (GBM112 and GBM144) yielded only the CTRL-GSC, while 14 cases did not generate any GSC (Fig. 1B; Table EV1).

Among all cases where at least the CTRL-GSC was established (n=11), the GSC derivation efficiency was 44%, which is lower than the 61% efficiency observed under similar culture conditions in our previous study (De Bacco *et al*., 2023b). This difference may be due to the use of a conventional, rather than ultrasonic, aspirator for GBM surgical removal in the current study, potentially reducing cell viability and, consequently, GSC derivation.

In summary, we established a protocol for generating *ex vivo* models of GSCs that emerge following standard therapies. By combining clinically relevant therapeutic pressures, culture conditions that select for GSCs, and a brain-mimicking microenvironment enriched with survival-promoting growth factors, these models aim to recapitulate the evolution from de novo to recurrent GBMs and facilitate the dissection of the selective effects of chemotherapy and radiotherapy.

### GSC derivation under therapeutic pressure associates with AKT pathway genetic alterations in original GBMs

Next-generation sequencing (NGS) of the original GBMs (whole tumor cell populations), was performed using a targeted panel covering GBM driver genes, and DNA damage-repair genes (GBM-target panel) (De Bacco *et al*., 2023b). This analysis confirmed the absence of *IDH1/2* gene alterations (Fig. 1B). Consistent with previous reports (Killela *et al*, 2013), *TERT* promoter (p*TERT*) mutations were frequent (88%) (Fig. 1B). Because of its prevalence, p*TERT* variant allele frequency (VAF) was used to estimate tumor cell content in the original suspension (2 x VAF, range 0.24-64.88%) (Table EV2) (De Bacco *et al*., 2023b; Schulze Heuling *et al*, 2017). No association was found between higher tumor cell content (>10%) and the ability to derive res-GSC families (Fisher’s exact test, p=0.65).

In our cohort, frequencies of other driver alterations were comparable to those reported by TCGA(Brennan *et al*, 2013) and the GLASS consortium (Barthel *et al*, 2019), except for *CDKN2A/B* deletions, which may be underdetected due to low tumor cell content limiting copy-loss assessment (De Bacco *et al*., 2023b) (Fig. 1B; Table EV2).

When we tested associations between genetic alterations and res-GSC family derivation (n=9/25 GBMs), we found a significantly higher frequency of pathogenic alterations in AKT pathway genes, including *PTEN* and *PIK3CA*, in GBMs yielding GSC families (Fisher’s exact test, p=0.03) (Figs. 1B-C and EV1G; Table EV2). Given the role of AKT pathway in suppressing apoptosis, these variants may enhance GSC derivation under therapeutic pressure (Liu *et al*, 2020). We also observed a significant association between *TP53* mutations and res-GSC family derivation (Fisher’s exact test, p=0.0005), and a trend toward an inverse association with EGFR alterations (Figs. 1B-C and EV1G). However, our previous study showed that *TP53* mutations and *EGFR* amplification respectively promote or hinder GSC derivation *per se*, independent of therapeutic pressures (De Bacco *et al*., 2023b).

No other genetic alterations significantly correlated with res-GSC family generation (Figs. 1B-C and EV1G; Table EV2).

### MGMT expression in original GBMs predicts GSC derivation under temozolomide pressure

Next, we asked whether res-GSC family generation correlates with expression – in original GBMs - of O^6^-Methylguanine-DNA Methyltransferase (MGMT), an enzyme that repairs TMZ-induced DNA damage and is the main determinant of primary TMZ resistance (Fu *et al*, 2012). A clinical surrogate marker for MGMT expression is the methylation status of the *MGMT* promoter (Hegi *et al*, 2005): GBMs with a methylated *MGMT* promoter are defined as MGMT-neg, whereas those with unmethylated *MGMT* promoter are defined as MGMT-pos. Although no correlation was found between *MGMT* status and overall derivation of GSC families (Fig. EV1F), in 6 out of 7 MGMT-pos GBMs yielding at least the CTRL-GSC, we could derive the TMZ-GSC as well (Fig. 1D). As expected, ex-vivo TMZ treatment of the MGMT-pos GBM whole-cell populations did not significantly impact viability, allowing TMZ-GSCs to stabilize with no statistically significant difference in the derivation time (i.e., reaching p5) between paired TMZ-GSCs and CTRL-GSCs (Figs. 1D and EV2A). However, in one case (GBM95), the derivation time for the TMZ-GSC was 80% longer than for its matched CTRL-GSC, suggesting incomplete TMZ resistance (Figs. 1D and EV2A). Consistent with the unmethylated *MGMT* promoter status of their parental GBMs, all these TMZ-GSCs expressed MGMT protein, as did their matched CTRL-GSCs (and IR-GSCs) (Fig. EV2B). The sole notable exception was MGMT-pos GBM149, which yielded a family whose members lacked MGMT protein expression despite retaining *MGMT* promoter unmethylation (Fig. EV2B; Table EV3). The hypothesis that this discrepancy was due to *MGMT* gene translocation (Oldrini *et al*, 2020) was ruled out (Fig. EV2C), leaving open the possibility of alternative epigenetic dysregulations (Hegi *et al*., 2005).

Concerning the remaining GBMs that yielded CTRL-GSCs (n=4), two were overtly MGMT-neg and two were *bona fide* MGMT-neg because, although *MGMT* status in the GBMs was undetermined due to insufficient material, the CTRL-GSCs were MGMT-neg (Fig. 1D; Table EV3). In these GBMs, TMZ significantly reduced viability of the initial cell population, resulting in generation of only two TMZ-GSCs with a two- or three-fold increase in derivation time compared to CTRL-GSCs (Figs. 1D and EV2A). In these families, all TMZ-GSCs (and IR-GSCs) lacked MGMT expression, as did CTRL-GSCs (Fig. EV2B).

Overall, we conclude that MGMT expression status of the original GBM influenced the derivation of TMZ-GSCs and that, whether positive or negative, MGMT status was preserved during propagation of the GSC family members, possibly due to inheritance of the underlying epigenetic trait.

Concerning derivation under IR pressure, we were able to derive IR-GSCs from 8 out of 11 GBMs that yielded at least the CTRL-GSC. In most cases, radiation modestly affected the viability of the initial cell population, resulting in only a modest increase in IR-GSC derivation time compared with CTRL-GSCs (median derivation-time increase of 14% vs. CTRL-GSCs, with a statistically significant difference between groups). The exception was the GBM95 family, in which IR was more effective on the initial cell population, and the IR-GSC was derived with a time increase of 121% relative to the CTRL-GSC (Figs. 1E and EV2A).

### Acquired resistance and drug tolerance to temozolomide in treatment-selected GSCs

In established res-GSC families, we investigated the TMZ response evolution by challenging treatment-naïve (CTRL-) GSCs as a surrogate of the response within the primary GBM (primary response), and treatment-selected (TMZ-) GSCs as a surrogate of the response within the relapsing GBM (secondary response) (Fig. 2A-K).

**Fig. 2.**
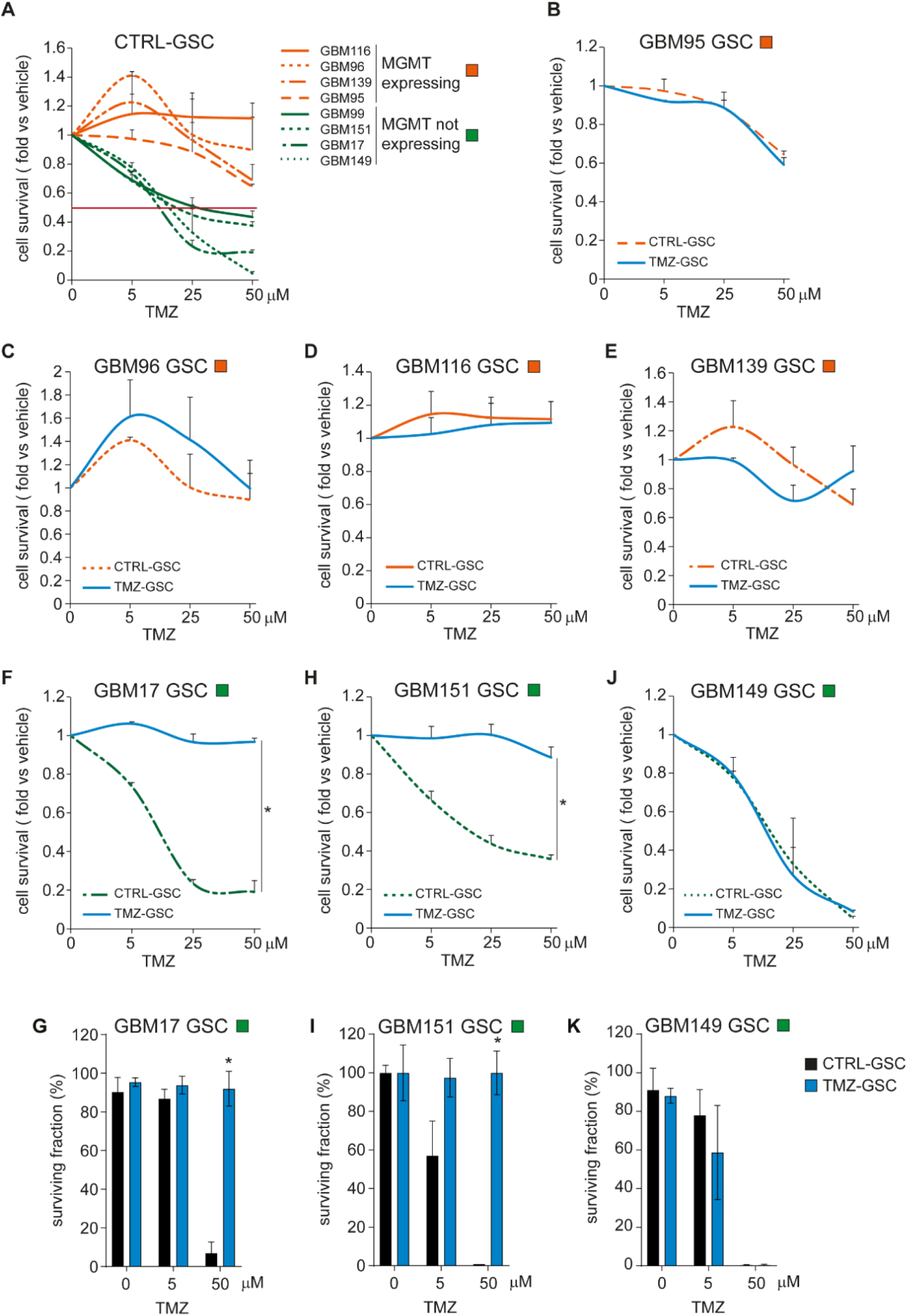
Acquired resistance and drug tolerance to temozolomide in treatment-selected GSCs. (A) Cell survival (fold vs. vehicle) of the indicated CTRL-GSCs, measured 9 days after TMZ treatment (5-50 µM). n = 2 independent experiments. (**B-F, H, J**) Cell survival (fold vs. vehicle) of matched CTRL-GSCs (as in A) and TMZ-GSCs of the indicated MGMT expressing (orange squares) or not expressing (green squares) res-GSC families, measured 9 days after TMZ treatment (5-50 µM). n = 2 independent experiments, Student’s paired t test. GBM17: *p=0.006256; GBM151: *p=0.013988. (**G, I, K**) Surviving fraction (%) of matched CTRL-GSCs and TMZ-GSCs from MGMT-non-expressing families (green squares), measured by clonogenic assay. n = 2 independent experiments, Student’s paired t test. GBM17: *p=0.000173; GBM151: *p=0.002529. In (A-K), data are presented as mean ± SEM.

In dose-response experiments, as expected, MGMT-expressing CTRL-GSCs exhibited primary resistance to the TMZ schedule used for selection (50 μM every three days for three cycles), i.e. they displayed cell survival >50% relative to vehicle-treated controls. In contrast, CTRL-GSCs lacking MGMT showed primary TMZ sensitivity (Fig. 2A).

Concerning the secondary TMZ response, TMZ-GSCs derived from MGMT-pos GBMs and expressing MGMT displayed resistance comparable to matched CTRL-GSCs, consistent with maintenance of primary resistance (Fig. 2B-E). Conversely, the two TMZ-GSCs selected from two MGMT-neg GBMs (GBM17 and GBM151), while still lacking MGMT expression (Fig. EV2B), developed secondary TMZ resistance compared to their matched, TMZ-sensitive CTRL-GSCs, as shown by survival curves and long-term clonogenic assays (Franken *et al*, 2006) (Fig. 2F-I). A distinct case was the GBM149 family, in which the TMZ-GSC remained highly TMZ sensitive, like its matched CTRL-GSC (Fig. 2J-K). However, unlike the other MGMT-neg families, GBM149 TMZ-GSC displayed a derivation time similar to CTRL-GSC (Figs. 1D and EV2B). The ability to survive selection while retaining TMZ sensitivity is consistent with a drug-tolerant persister phenotype (Pu *et al*, 2023; Russo *et al*, 2024), rather than development of stable secondary resistance.

In summary, in GSCs representative of the primary GBM (CTRL-GSCs), MGMT protein expression correlates with primary TMZ resistance, whereas its absence correlates with primary sensitivity. In GSCs derived under TMZ pressure (TMZ-GSCs), representative of the relapsing tumor, we observed either evolution from primary sensitivity to secondary resistance or drug tolerance, neither associated with MGMT upregulation.

### Genetic alterations driving secondary resistance to temozolomide in MGMT-neg GSCs

To investigate mechanisms underlying secondary TMZ resistance or tolerance, we considered the emergence of GSC subclones harboring genetic alteration(s), either pre-existing in the treatment-naive GBM or induced by ex vivo treatment, which could favor subclone survival under therapeutic pressure (Fu *et al*., 2012; Woo *et al*, 2019). We therefore compared the genetic landscape of MGMT-neg TMZ-GSCs (GBM17, GBM151, and GBM149) with their matched CTRL-GSCs. Using the GBM-target panel (De Bacco *et al*., 2023b), we found that typical GBM driver alterations, such as those affecting the *TERT* promoter, *PTEN* and *TP53* – already detectable in the original GBM cell suspension (Fig. 1B; Table EV2) - were retained across res-GSC family members, with VAFs consistent with GSC monoclonality (Fig. 3A; Table EV4). In contrast, EGFR amplification detectable in the original GBM17 tumor was lost in GSC family members, likely due to culture conditions (De Bacco *et al*., 2023b; Schulte *et al*, 2012) (Fig. 3A; Tables EV2 and EV4).

**Fig. 3.**
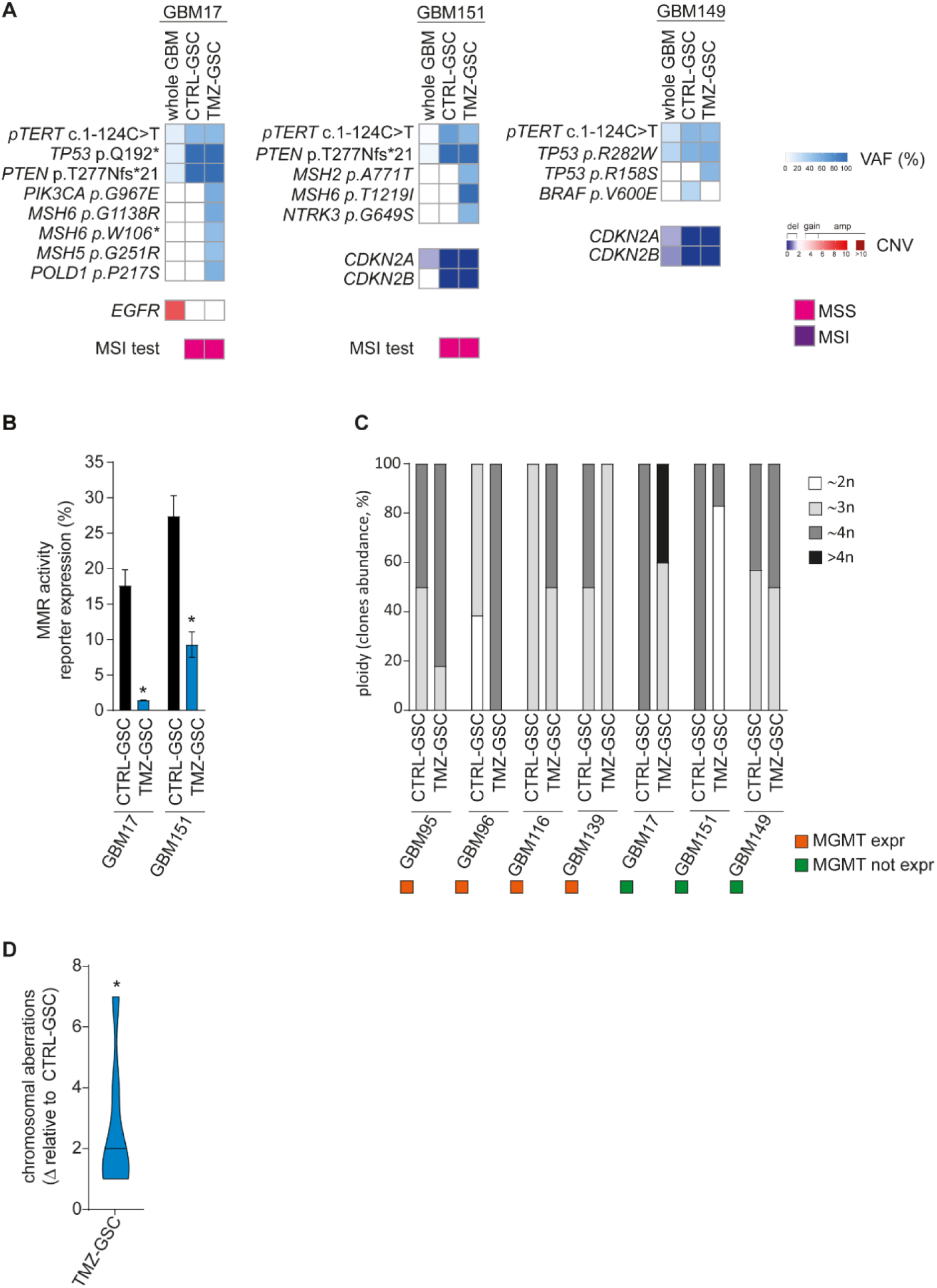
Genetic alterations driving secondary resistance to temozolomide in MGMT-neg GSCs. (**A**) Pathogenic alterations (variant allele frequency, VAF, and gene copy number variation, CNV) detected by the NGS GBM-target panel or ddPCR (p*TERT*) in MGMT-neg whole GBMs and matched CTRL-GSCs and TMZ-GSCs. MSS: microsatellite stable; MSI: microsatellite instable, assessed at passage 10. (**B**) MMR activity measured by FM-HCR in matched CTRL-GSC and TMZ-GSC from GBM17 and GBM151 families, 24 hours after transfection. Data are z-values (mean ± SEM). n = 2 independent experiments for GM17 and n = 3 independent experiments for GBM151, Student’s paired t test. GBM17: *p=0.018734; GBM151: *p= 0.006080. (**C**) Ploidy (n) in MGMT-pos (orange square) and MGMT-neg (green square) TMZ-GSCs and CTRL-GSCs assessed by chromosome G-banding. (**D**) Cumulative chromosomal aberration number in all TMZ-GSCs vs. matched CTRL-GSCs, assessed by G-banding and multicolor fluorescent in situ hybridization (M-FISH). Data are shown as delta versus the corresponding CTRL-GSC; median is indicated. One sample Wilcoxon test. *p=0.0156.

In secondary resistant MGMT-neg TMZ-GSCs (GBM17 and GBM151), we observed the emergence of private pathogenic alterations that were undetectable in both the original GBM population and the matched CTRL-GSC (Fig. 3A; Tables EV2 and EV4). These included mutations of *MSH2* and *MSH6*, key components of the DNA Mismatch Repair (MMR) machinery, whose deficiency confers TMZ resistance by preventing futile DNA repair cycles and subsequent apoptosis (Fig. 3A; Table EV4) (Fu *et al*., 2012; Hickman & Samson, 2004; Li *et al*, 2025). Specifically, GBM17 TMZ-GSC harbored *bona fide MSH6* biallelic inactivation, whereas GBM151 TMZ-GSC harbored *MSH6* biallelic and *MSH2* monoallelic inactivation (Fig. 3A; Table EV4). These alterations functionally impaired MMR, as shown by a fluorescence-based multiplex flow-cytometric host cell reactivation assay (FM-HCR), which uses a fluorescent reporter detected only in MMR-proficient cells (Fig. 3B) (Nagel *et al*, 2014; Piett *et al*, 2021). Both *MSH6* variants (p.T1219I and p.G1138R) have been reported in association with TMZ treatment (Crisafulli *et al*, 2022). These alterations did not induce a shift towards microsatellite instability (MSI) at GSC establishment (p10) or at later passages (Fig. 3A), consistently with reported outcomes of biallelic *MSH6* or monoallelic *MSH2* inactivation (Helderman *et al*, 2025; Kets *et al*, 2006; Salem *et al*, 2020). The mutation spectrum (predominantly C>T/G>A; Table EV4) supports de novo TMZ mutagenesis followed by selection of the MMR-deficient, TMZ-resistant clones, consistent with findings in GBM tissue studies (Barthel *et al*., 2019; Touat *et al*, 2020) and matched GSCs from primary and recurrent GBMs (Orzan *et al*, 2017). Additional TMZ-associated mutations detected in TMZ-GSCs, while likely pathogenic, affected genes (including *PIK3CA* and *NTRK3*) unlikely to mediate robust TMZ resistance mechanisms but that could contribute to tumor evolution (Fig. 3A; Table EV4).

In the GBM149 TMZ-GSC, which appeared to tolerate TMZ during selection process yet remained TMZ sensitive afterward, no MMR alteration emerged. Instead, comparison with CTRL-GSC revealed an additional *TP53* mutation, unlikely to reflect TMZ mutagenesis (Fig. 3A; Table EV4). Conversely, a *BRAF* V600E mutation was detected in CTRL-GSC but not in TMZ-GSC (Fig. 3A). Both remained undetected in the original GBM population, leaving unresolved whether pre-existing subclones were selected during GSC propagation (Fig. EV3A). Notably, the *TP53* mutation present in the original GBM may have promoted subclonal heterogeneity (Figs. 3A and EV3A; Table EV4).

As expected, MGMT-pos TMZ-GSCs shared driver alterations with matched CTRL-GSCs, with no additional private alterations except in GBM102 TMZ-GSC (Table EV4).

The broader impact of TMZ treatment was assessed by cytogenetics, which showed marked karyotypic changes, including ploidy shifts and chromosomal aberrations, in both MGMT-neg and MGMT-pos TMZ-GSCs relative to CTRL-GSCs (Figs. 3C-D and EV3B; Tables EV4 and EV5).

In summary, MGMT-neg TMZ-GSCs with secondary resistance are genetically distinct from their treatment-naïve counterparts, consistent with TMZ mutagenesis combined with selection for MMR inactivation, which confers stable resistance. The model also captures other TMZ response modes, including drug tolerance in MGMT-neg GSCs lacking MMR mutations, and substantial cytogenetic remodeling even in primary resistant (MGMT-pos) settings.

### Evolution of radioresistance in GSCs under IR pressure

Next, we investigated radioresistance evolution by challenging CTRL-GSCs and IR-GSCs as surrogates of the response within the primary GBM (primary response) and the relapsing GBM (secondary response), respectively (Figs. 4A-C and EV4A-B).

**Fig. 4.**
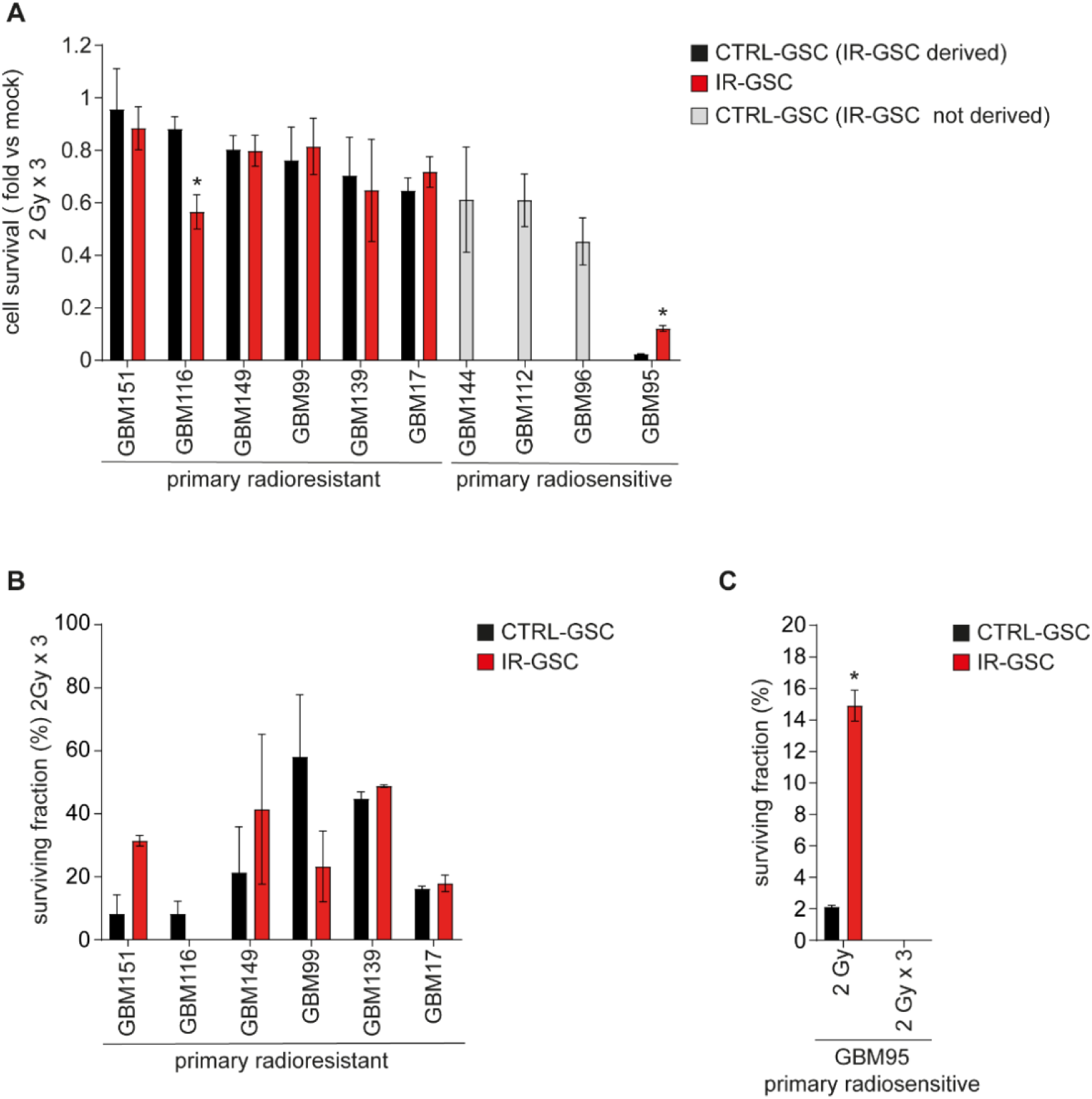
Evolution of radioresistance in GSCs under ionizing radiation pressure. (**A**) Cell survival (fold vs. mock) of matched CTRL-GSCs and IR-GSCs irradiated with 2 Gy x 3 days, measured 48 h after the last dose. Mock: no IR. n = 2 independent experiments for all cases but GBM99 (n = 3 independent experiments), Student’s paired t test. GBM116: *p= 0.023240, GBM95: *p=0.013187. (**B**) Surviving fraction (%) of primary radioresistant CTRL-GSCs and matched IR-GSCs, measured by clonogenic assay after irradiation (2 Gy for 3 days). n = 2 independent experiments, Student’s paired t test. (**C**) Surviving fraction (%) of GBM95 CTRL-GSC and IR-GSC, measured by clonogenic assay after a single 2 Gy dose or 2 Gy x 3 days. n = 2 independent experiments, Student’s paired t test. *p= 0.000104. In (A-C), data are presented as mean ± SEM.

After treating CTRL-GSCs with 2 Gy for three consecutive days - a dose that discriminates resistant from sensitive GSCs (De Bacco *et al*., 2016) - 6 out of 10 GSCs were primarily radioresistant (cell survival >60% vs. untreated), whereas 4 out of 10 were radiosensitive, suggesting that their parental GBMs were radioresistant or radiosensitive, respectively (Fig. 4A).

From the six primary radioresistant GBMs, IR-GSCs were derived in all cases (Fig. 4A). When challenged to assess the secondary response (2 Gy x 3 days), these IR-GSCs showed no significant differences relative to matched CTRL-GSCs, in cell survival or in more stringent radiobiological clonogenic assays (Fig. 4A-B). An exception was GBM116 IR-GSC, which displayed reduced radioresistance in cell survival assays (Fig. 4A). Similar results were obtained with a single 5 Gy dose, biologically equivalent to 2 Gy x 3 (Fig. EV4A-B).

From the four primary radiosensitive GBMs, only one IR-GSC could be derived (GBM95) (Fig. 4A). GBM95 IR-GSC showed a significant, though incomplete, increase in radioresistance compared to its matched CTRL-GSCs. This was detected in cell survival assays after treatment with 2 Gy x 3 days or 5 Gy (Figs. 4A and EV4A), and in radiobiological clonogenic assays with a single 2 Gy dose (Fig. 4C). We therefore classified GBM95 IR-GSC as secondary radioresistant.

In summary: (i) more than half of GBMs display high primary radioresistance (as reflected by CTRL-GSCs), and yield IR-GSCs that maintain this phenotype; (ii) primary radiosensitivity limits IR-GSC derivation under IR pressure; and (iii) when IR-GSCs can be derived from radiosensitive GBMs, they can evolve secondary (albeit partial) radioresistance.

### Secondary GSC radioresistance is associated with enhanced DNA damage response and cell cycle adaptation

As with TMZ treatment, we hypothesized that IR therapeutic pressure could select pre-existing, or induce *de novo*, genetic alterations that promote radioresistance and tumor progression. We therefore evaluated the genetic features of the secondary radioresistant GBM95 IR-GSC, and of IR-GSCs derived from primary radioresistant GBMs. GBM-target panel analysis revealed that all IR-GSCs, including the secondary radioresistant, shared the same alterations as their matched CTRL-GSCs, with the sole exception of GBM149 family, in which, as mentioned above, CTRL-GSCs harbored a BRAF mutation that was absent in the matched IR-GSC (Fig. 5A; Table EV4) and TMZ-GSC (Fig. 3A).

**Fig. 5.**
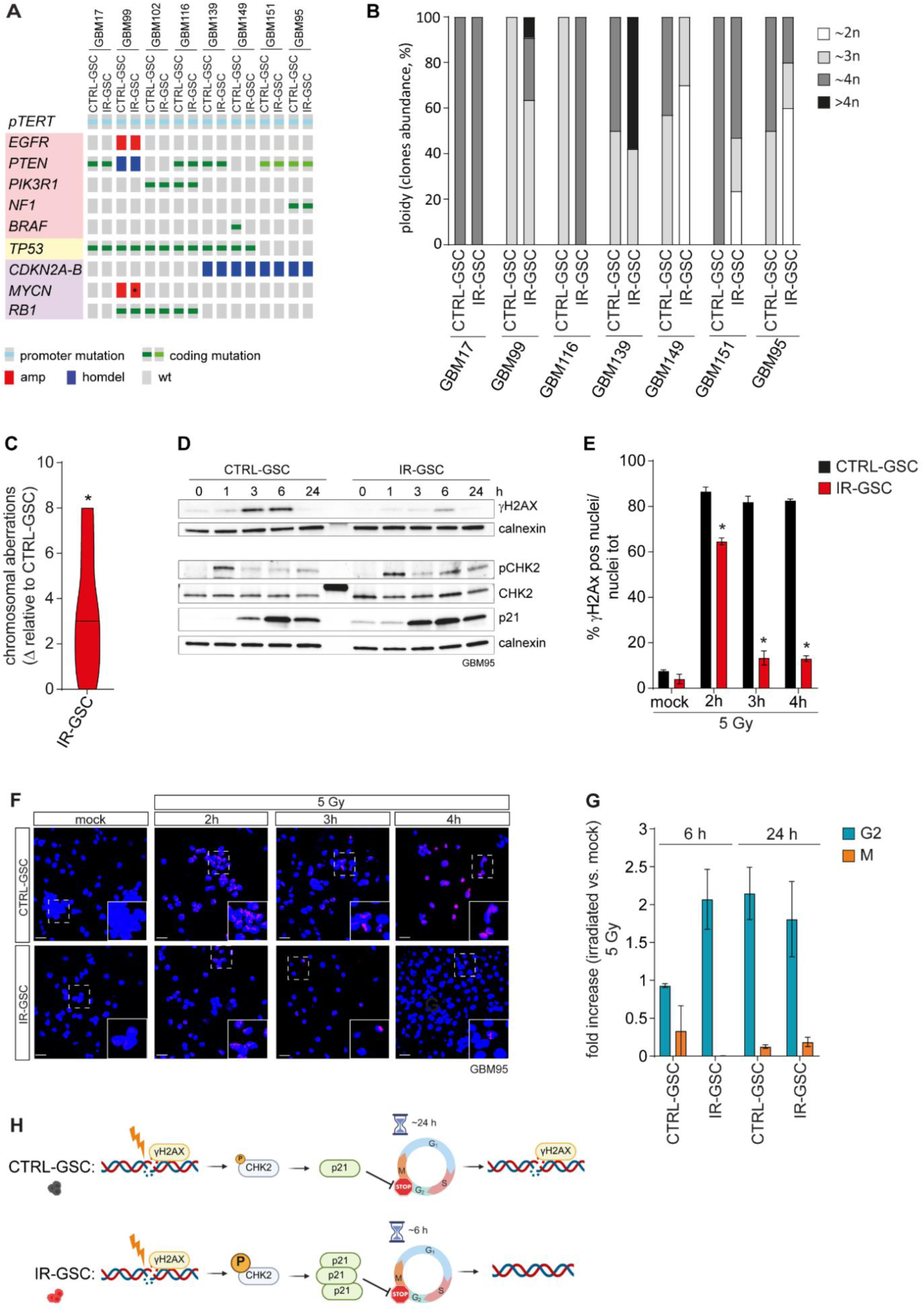
Secondary GSC radioresistance is driven by enhanced DNA damage response and cell cycle adaptation. (**A**) Genetic profiling of matched CTRL-GSCs and IR-GSCs using the NGS GBM-target panel; p*TERT* mutations were assessed by ddPCR. Samples are ordered by primary and secondary radioresistance. *Significant increase in *MYCN* copy number; see also A Table EV4. (**B**) Ploidy (n) of matched CTRL-GSCs e IR-GSCs assessed by chromosome G-banding. (**C**) Cumulative chromosomal aberration number in all IR-GSCs vs. matched CTRL-GSCs, assessed by G-banding and M-FISH. Median number is indicated. Data are shown as delta relative to the corresponding CTRL-GSC; median is indicated. One-sample Wilcoxon test. *p = 0.0313. (**D**) Western blots showing phospho-H2AX (γH2AX), phospho-CHK2 (pCHK2; Tyr68), total CHK2 and p21 in GBM95 CTRL-GSC and IR-GSC in a representative time-course experiment after a single 5 Gy dose. Calnexin: loading control. (**E-F**) Immunofluorescence for phospho-H2AX (γH2AX) in GBM95 CTRL-GSC and IR-GSC in a time-course experiment after a single 5 Gy dose. (**E**) Quantification of γH2AX-positive cell nuclei. n = 4 technical replicates, Student’s paired t test. * p<0.000001. (**F**) Representative images. Scale bar, 25 μm. (**G**) Fraction of GBM95 CTRL-GSC and IR-GSC cells in G2 and M phases, measured by flow cytometry using phospho-Histone 3 (pH3; Ser10) staining at 6 and 24 h after a single 5 Gy dose, shown as fold change vs. mock. Mock: no IR. n = 2 independent experiments, Student’s paired t test. (**H**) Schematic of adaptive mechanisms of secondary radioresistance in GBM95 GSC family. In (E, G), data are presented as mean ± SEM.

In contrast, all IR-GSCs showed karyotypic changes, including ploidy variation and a statistically increased number of chromosomal aberrations compared with CTRL-GSCs, similar to TMZ-GSCs (Fig. 5B-C and EV3B; Table EV5).

Because radioresistance can also rely on adaptive mechanisms that enhance the DNA Damage Response (DDR) (Ahmed *et al*, 2015; Bao *et al*., 2006; De Bacco *et al*., 2016; Squatrito & Holland, 2011), we assessed the ability of the secondary radioresistant GBM95 IR-GSC to repair radiation-induced DNA damage. In time-course experiments, this IR-GSC showed a stronger DDR than CTRL-GSC. Specifically, it exhibited (i) increased DDR activation, indicated by sustained CHK2 phosphorylation, and (ii) more efficient DNA damage resolution, indicated by reduced phosphorylation of histone H2AX (γH2AX), a marker of DNA double-strand breaks, in western blot and immunofluorescence (Fig. 5D-F). No significant changes in the protein levels of the DNA damage sensor MRN complex (including MRE11, RAD50 and NBS1) or in RAD51, a key effector of DNA homologous recombination (HR), were observed (Fig. EV4C-D).

IR also induced the cyclin-CDK complex inhibitor p21 (a p53 target) more strongly in IR-GSC than in CTRL-GSC at an early time-point (Fig. 5D). This increased p21 induction was associated with a more effective G2/M block, with cell accumulation in G2 and absence of cells in M phase 6 h after irradiation (Fig. 5G). In contrast, CTRL-GSC still progressed through G2/M at 6 h and reached a G2/M distribution comparable to IR-GSC only at 24 h.

In summary, (i) IR-selected GSCs accumulate chromosomal-level alterations regardless of primary radioresistance, and (ii) secondary radioresistance is associated with enhanced double-strand break repair, consistent with adaptive DDR activation coordinated with G2/M arrest (Fig. 5H).

### GSCs emerging from treatments exhibit marker shifts despite stable transcriptional subtypes

Therapeutic pressure is known to modify the GBM transcriptional profile, with shifts towards mesenchymal, neuronal and proliferative phenotypes (Garofano *et al*, 2021; Hoogstrate *et al*, 2023; Kim *et al*, 2024; Liu *et al*, 2024; Tanner *et al*, 2024; Varn *et al*, 2022; Wang *et al*, 2022; Wang *et al*, 2017).

To assess transcriptional changes in the res-GSC family members, we applied the established signatures from Wang (Wang *et al*., 2017) and Neftel (Neftel *et al*, 2019) (descriptive criteria) and from Garofano (Garofano *et al*., 2021) (metabolic/functional classification). Wang and Neftel subtyped GSCs consistently and revealed minimal within-family heterogeneity, except in GBM151, where the secondary resistant TMZ-GSC shifted toward a classical subtype by Wang’s signature only (Fig. 6A; Tables EV6 and EV7; Dataset EV1). Comparing the two methods, we note that: (i) mesenchymal assignments (5/9 families) overlapped one-to-one; (ii) among families subtyped as classical by Wang (3/9), two (GBM99 and GBM116) corresponded to Neftel’s astrocyte (AC)-like subtype, whereas in the GBM95 family, the AC-like subtype coexisted with neuronal progenitor cell (NPC)-like and oligodendroglial progenitor cell (OPC)-like profiles; notably, the secondary radioresistant GBM95 IR-GSC showed stronger activation of both Wang’s proneural signature and Neftel’s NPC2-like profile; and (iii) the GBM96 family, subtyped as proneural by Wang, coherently showed NPC-like and OPC-like profiles, along with an AC-like profile, according to Neftel (Fig. 6A; Tables EV6 and EV7; Dataset EV1).

**Fig. 6.**
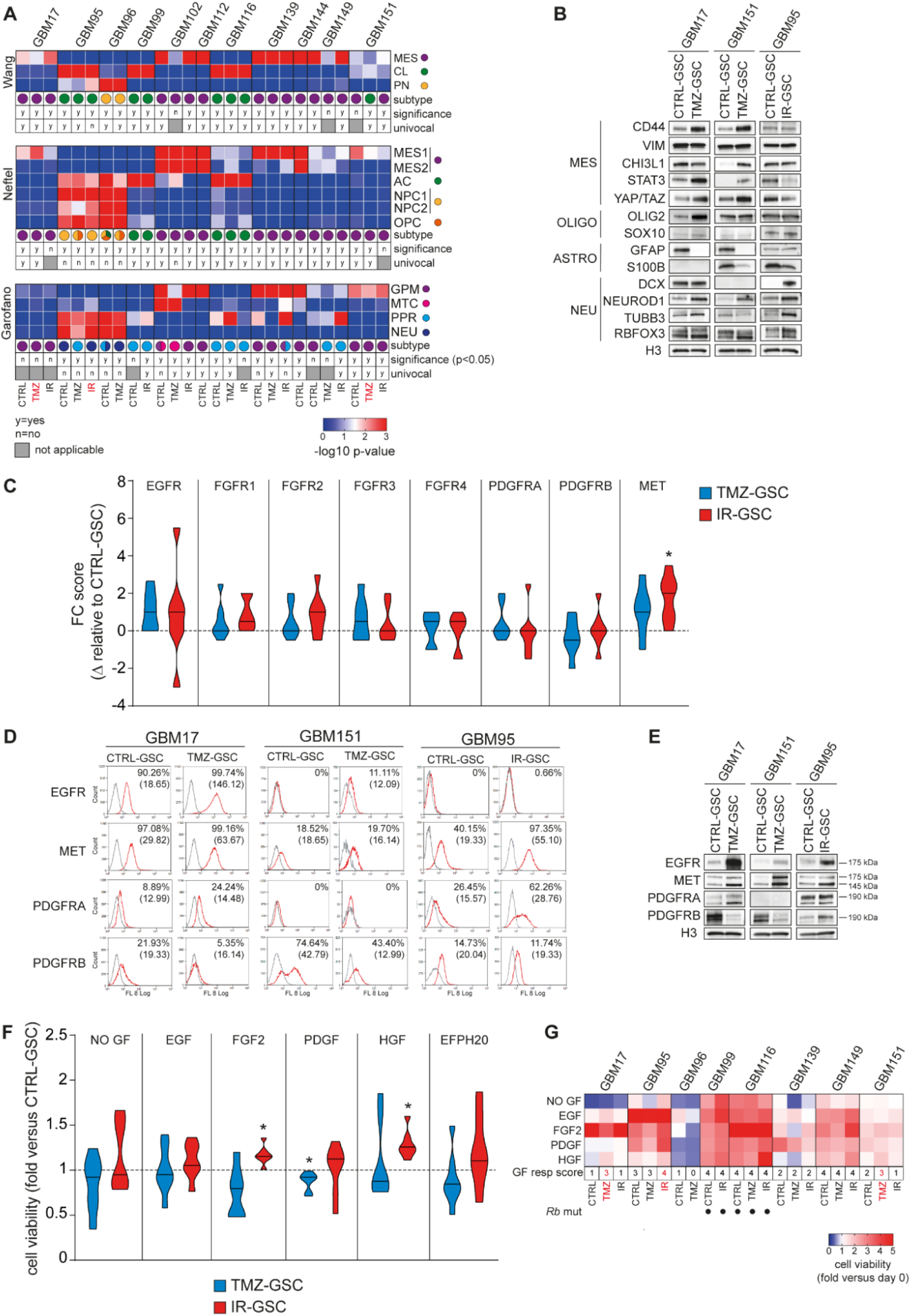
GSCs emerging from treatments exhibit phenotypic marker shifts and increased RTK expression and activity. (**A**) Transcriptional classification of res-GSCs according to Wang (MES, mesenchymal; CL, classical; PN, proneural), Neftel (MES1/2, mesenchymal-like 1/2; AC, Astrocyte-like; NPC1/2, neural progenitor-like 1/2; OPC, oligodendrocyte progenitor-like), and Garofano (GPM, glycolytic/plurimetabolic; MTC, mitochondrial; PPR, proliferative/progenitor; NEU, neuronal) signatures. Secondary resistant GSCs are indicated in red. Color intensity is proportional to subtype-assignment probability. Subtype assignment is based on the lowest p-value (significant or not). Univocal: only one significant subtype assignment. (**B**) Western blots showing mesenchymal and lineage markers in GBM17, GBM151 and GBM95 CTRL-GSCs and matched secondary resistant family members. H3: loading control. MES, mesenchymal; OLIGO, oligodendroglial; ASTRO, astrocytic; NEU, neuronal markers. (**C**) Flow-cytometry (FC) score of the indicated RTK surface expression grouped by family member. Scores combine percent positive cells and median fluorescence intensity (see Methods). Data are shown as delta (Δ) relative to the matched CTRL-GSCs; median is indicated. One-sample Wilcoxon test. *p= 0.0313. (**D**) Flow-cytometry analysis of surface expression of the indicated RTKs in secondary resistant GSCs and matched CTRL-GSCs (GBM17, GBM151 and GBM95). Percent positive cells and mean fluorescence intensity are shown. (**E**) Western blots showing expression of the indicated RTKs in secondary resistant GSCs and matched CTRL-GSCs (GBM17, GBM151 and GBM95). H3: loading control. Molecular weights are indicated. (**F**) Cell viability (normalized versus day 0) of TMZ- and IR-GSCs, measured in the presence or absence of the indicated growth factors. Data are shown as fold change relative to the matched CTRL-GSCs. One sample Wilcoxon test. *p= 0.0156. (**G**) Heatmap of cell viability (normalized versus day 0) for all GSCs in the presence or absence of the indicated growth factors. Growth responsiveness (GF resp) score is shown.Secondary resistant GSCs are indicated in red. Dots indicate the presence of *RB1* mutations.

Subtyping according to Garofano’s signatures showed strong correspondence between the glycolytic/plurimetabolic (GPM) subtype and a mesenchymal profile, consistent with prior findings (Garofano *et al*., 2021), and between the neuronal (NEU) subtype – often co-occurring with the proliferative/progenitor (PPR) profile - and NPC-like states (Fig. 6A; Tables EV6, EV7 and EV8; Dataset EV1). This approach revealed greater within-family heterogeneity. In particular, PPR scores were increased in several therapy-emergent members, independent of secondary resistance (GBM99 IR-GSC; GBM116 TMZ-GSC; GBM139 IR-GSC and GBM149 IR-GSC) (Fig. 6A; Tables EV6, EV7 and EV8; Dataset EV1).

Although global subtyping showed limited within-family change, analysis of subtype markers revealed clearer phenotypic shifts in secondary resistant GSCs. Secondary resistant TMZ-GSCs (GBM17 and GBM151) showed upregulation of mesenchymal (Carro *et al*, 2010; Kim *et al*, 2021; Verhaak *et al*, 2010) and neural-lineage markers (Pagani *et al*, 2025; Verhaak *et al*., 2010) with downregulation of the astrocytic marker GFAP, whereas the secondary radioresistant GBM95 IR-GSC showed reduced mesenchymal markers along with increased neuronal-lineage markers (Fig. 6B).

In summary, treatment-emergent GSCs show limited changes in commonly used GBM transcriptional profiles, as compared with matched CTRL-GSCs, which are representative of treatment-naive GSCs residing in the primary GBM. However, treatment-emergent GSCs can upregulate the PPR program, which includes genes linked to proliferation and DNA repair, and GSCs becoming secondary resistant display marker-level shifts toward mesenchymal or neuronal phenotypes.

### GSCs emerging from treatments exhibit increased RTK expression and growth factor responsiveness

Chemo- and radiotherapy pressures can promote GSC resistance through hyperactivation of survival pathways sustained by increased expression and activation of Receptor Tyrosine Kinases (RTKs) (De Bacco *et al*., 2016; Gimple *et al*., 2022; Squatrito & Holland, 2011). Flowcytometric analysis of cell-surface RTKs commonly expressed (and/or genetically altered) in GBM (EGFR, FGFR1-4, PDGFRA-B and MET) (Brennan *et al*., 2013; Ozawa *et al*, 2010; Szerlip *et al*, 2012) showed a trend toward increased expression in res-GSC family members emerging under therapeutic pressure across families, particularly in IR-GSCs, irrespective of resistance acquisition or increase. This trend reached statistical significance for MET, consistent with its established role in promoting radioresistance (De Bacco *et al*., 2016; De Bacco *et al*, 2011; Joo *et al*, 2012) (Figs. 6C and EV5A).

Detailed analysis revealed that EGFR, MET and PDGFRA (but not PDGFRB or FGFR 1-4) were significantly more expressed at the cell surface and/or in the cytoplasm of secondary resistant GSCs compared with CTRL-GSCs, with increased RTK expression at single cell level (indicated by mean fluorescence intensity) and an increased fraction of RTK-positive cells (Figs. 6D and EV5B-D). These changes were confirmed by total RTK protein analysis in western blot (Fig. 6E). RTK protein increases in secondary resistant GSCs were not associated with gene copy-number gains (Fig. EV3B and Table EV5) or higher mRNA expression (Dataset EV1) relative to CTRL-GSCs, suggesting the establishment of post-transcriptional regulation mechanisms.

Changes in RTK expression were reflected in proliferative and viability response to the corresponding growth factors (GF). Overall, IR-GSCs showed a trend toward increased proliferative rate and/or viability, reaching significance for responses to FGF2 and HGF, whereas TMZ-GSCs tended to reduced proliferation and/or viability, reaching significance for PDGF, compared to CTRL-GSCs (Fig. 6F). Detailed analysis showed that, consistent with specific RTK upregulation, secondary resistant GSCs displayed stronger responses to individual GFs, and a higher cumulative GF-response score, compared to CTRL-GSCs (Fig. 6G).

Finally, assessing GF deprivation showed that GSCs (either treatment-naïve or emerging from therapies) could proliferate autonomously only when harboring Rb mutations (GBM99 and GBM116) (Fig. 6G; Table EV4). In contrast, the other GSC families were unable to proliferate and/or to survive, indicating continued dependence on exogenous GFs (Fig. 6G).

In conclusion, GSCs emerging under therapeutic pressures - particularly those evolving secondary resistance- show increased RTK expression, with MET and EGFR playing prominent roles, and increased responsiveness to GFs required for their survival and propagation.

## DISCUSSION

Over the past decade, longitudinal profiling of GBM tissues has clarified key aspects of evolution from primary to recurrent tumor and linked genetic and adaptive programs to progressive therapy refractoriness (Barthel *et al*., 2019; Hoogstrate *et al*., 2023; Johnson *et al*, 2014; Kim *et al*, 2015; Kim *et al*., 2024; Kocakavuk *et al*, 2021; Lucas *et al*, 2025; Spitzer *et al*, 2025; Varn *et al*., 2022; Wang *et al*, 2016). Translation has nonetheless lagged, in part due to a lack of experimentally tractable systems that capture heterogeneity and evolution at the GSC level, a key driver of resistance and relapse (Gimple *et al*., 2022; Osuka & Van Meir, 2017; Sloan *et al*., 2024).

To address this, we developed an ex vivo derivation protocol to generate res-GSC families that approximate matched GSC states from primary and recurrent GBM. The platform is built on: (i) tumor collection by surgical aspirators to sample broadly and preserve heterogeneity; (ii) propagation in growth factor–enriched medium (EGF, FGF2, PDGF, HGF) that better reflects brain and GBM cues than the conventional stem medium containing EGF and FGF2 only (De Bacco *et al*., 2023a; De Bacco *et al*., 2023b); (iii) early application of therapeutic pressure soon after dissociation, limiting culture-driven drift and ensuring uniform exposure while cells remain largely as single elements; and (iv) clinically relevant TMZ and IR dosing schedules. Together, these elements create a controlled system in which therapy-naïve CTRL-GSCs and pressure-selected TMZ- and IR-GSCs can be derived from the same specimen, enabling within-patient, pressure-specific comparisons that are rarely feasible with matched primary/recurrent material.

Concerning the evolution of TMZ response, our results highlight the following aspects. First, the status of MGMT protein expression in the primary GBM, mainly determined by promoter methylation status (Hegi *et al*., 2005), plays a central role in conferring primary resistance or sensitivity, i.e., the response of the primary GBM at first drug administration. As known, MGMT protein activity confers primary resistance whereas lack of MGMT confers vulnerability to DNA alkylation induced by TMZ (Fu *et al*., 2012). Our results indicate that MGMT promoter methylation status is an epigenetic trait that appears to be stably inherited throughout multiple cell generations, as we observed by comparing the initial GBM cell populations and GSCs cultured for multiple passages. As also GSCs emerging from TMZ or IR therapeutic pressure continue to lack MGMT promoter methylation and express MGMT protein, we can infer that the same may occur in recurrent GBM, supporting the maintenance of TMZ resistance. In addition, our results highlight an unexpected finding: although primary resistant to TMZ (i.e., able to survive to the drug, almost behaving like matched controls not exposed to the drug), MGMT expressing GSCs are not fully shielded from clinically relevant doses of TMZ. Rather, they undergo genetic damage (DNA base alkylation) that they counteract with efficient MGMT-initiated DNA repair. However, this response does not lead to full *restitutio ad integrum*; instead, it results in broad chromosomal-level changes, which in principle could lead to translocations and aberrations endowing GSCs with selective advantages for growth in the harsh microenvironment induced by treatments (featuring inflammation, wound healing, DNA damage response etc.). This observation cautions against indiscriminate TMZ administration to GBM patients: in cases with MGMT expression, not only TMZ fails to provide a clinical benefit (Hegi *et al*, 2024), but it may be detrimental in perspective.

Second, in GBMs lacking MGMT expression, GSCs are primary sensitive to the drug, i.e. they undergo massive cell death together with the remaining GBM cell population. However, a GSC subpopulation can emerge under TMZ selective pressure, highlighting two possible scenarios of drug response evolution in the recurrent tumor, neither of which implies reactivation of MGMT expression. In the first scenario, GSCs undergo intense and widespread TMZ-driven mutagenic activity, followed by selection of subclone(s) harboring mutations in genes that provide the most efficient bypass of TMZ-induced cell death, namely the loss of MMR genes. Although this genetic alteration has been described as a mechanism of secondary TMZ resistance in recurrent GBM tissues and GSCs (Orzan *et al*., 2017; Touat *et al*., 2020), our model provides experimental evidence in a controlled system that MMR alterations associated with TMZ resistance disrupt MMR-mediated DNA repair, and offers a preclinical setting in which to assess much needed therapeutic strategies for recurrent GBM. Of note, the MSH6 loss preferentially induced and selected by TMZ treatment (which is present in both GSCs acquiring secondary TMZ resistance) do not elicit microsatellite instability, consistently with previous reports (Helderman *et al*., 2025; Kets *et al*., 2006; Salem *et al*., 2020). This may limit the efficacy of therapeutic interventions with drugs that proved to be effective in MMR-deficient tumors with microsatellite instability, such as WRN helicase inhibitors (Morales-Juarez & Jackson, 2022).

In a second scenario (GBM149, not expressing MGMT), TMZ caused, as expected, massive cell death; however this was followed by rapid recovery and emergence of GSCs that neither acquired MMR gene alteration nor reactivated MGMT expression and, accordingly, continued to remain TMZ sensitive. This TMZ-sensitive but resilient GSC family represents a model to study the drug tolerant, persister (DTP) phenotype, which remains poorly characterized in GBM and is better known in relation to therapies targeting cell proliferation drivers (Liau *et al*, 2017; Russo *et al*., 2024). Notably, the GSCs emerging from TMZ selective pressure harbored a TP53 mutation poorly compatible with TMZ mutagenesis. This mutation was absent from controls, indicating that a GSC subclone present in the original GBM, or that arose during cell propagation, was favored during the selection process. Even though the DTP state is believed to be mostly adaptive (Russo *et al*., 2024), these findings point to a role of TP53 inactivation in fostering specific aspects of the DTP state, especially those related to DNA damage tolerance, which is particularly relevant for TMZ treatment. This model could enable studies of the DTP state in GBM and therapeutic approaches that consider genotype and genotype-phenotype correlates of this DTP state.

A third aspect concerning the evolution of therapeutic response relates to IR. GSCs are known to be highly resistant to doses used in the clinics, compared with their pseudodifferentiated progeny; this radioresistance is attributed to the ability to promptly unleash the DNA damage response through kinases such as ATM and CHK2, activated by double-strand breaks and further stimulated by RTKs, in particular MET (Bao *et al*., 2006; De Bacco *et al*., 2016). Nevertheless, in our models, as well as in an independent GSC panel (De Bacco *et al*., 2016), differences in response to radiation - i.e., primary resistance vs. primary sensitivity – were evident, and influenced the ability to derive IR-selected cultures. In this respect, the res-GSC platform helps disentangle two clinically relevant situations: (i) tumors whose stem-like compartment is intrinsically radioresistant (and therefore tends to persist through IR without necessarily “needing” additional resistance evolution), and (ii) tumors that are initially radiosensitive, where selection is harsher and emergence of IR-surviving GSCs is less frequent, but can reveal adaptive routes to increased resistance.

In all cases of primary resistance, IR-GSCs did not display a further increase in radioresistance. In the case of the radiosensitive GBM95, a significantly increased, although incomplete, secondary resistance developed, allowing us to gain insights into the evolution of the IR response. All IR-GSCs, whether primary or secondary radioresistant did not display mutations or copy number variations in driver genes or DNA repair genes, as compared with matched CTRL-GSCs. Although the gene panel analyzed is limited, this lack of alterations was expected based on evidence that, in patients, radiotherapy is not associated with a significant increase in SNV burden, but rather with a DSB repair scar dominated by deletions, usually resulting from Non-Homologous Repair End-Joining (NHEJ) (Kocakavuk *et al*., 2021). However, in all IR-GSCs, chromosomal-level alterations accumulated compared to CTRL-GSCs, as result of DNA damage, and/or selection of subclones with increased genetic alterations. Whether chromosomal rearrangements could lead to fusion genes fostering radioresistance warrants further investigation. While information on gene sequencing is ample, the study of chromosomal changes in GBM recurrence are limited, mostly performed on whole tissues and, with notable exceptions (Kocakavuk *et al*., 2021), without distinguishing between effects of chemo- and radiotherapy (Zhang *et al*, 2023). Therefore, res-GSC models are uniquely suited to further pursue this aspect in a controlled, pressure-specific manner.

A further aspect emerging from the study of res-GSC families is that, in the evolution of therapeutic response - either to TMZ or IR- adaptation supported by transcriptional (or post-transcriptional) changes may play a major role. This is evident in the secondary resistant IR-GSC, where a more efficient execution of DNA damage repair was associated with increased activity of proteins responsible for initiating the DNA damage response. Transcriptomic profiles used for GBM subtyping tended to remain stable in GSC emerging from therapeutic pressures compared to controls, without overt shifts toward the ‘mesenchymal’ or ‘neuronal’ profiles described in tissue (bulk or single-cell) studies (Kim *et al*., 2024; Varn *et al*., 2022; Wang *et al*., 2022). Because the mesenchymal subtype is also clearly recognized in purified GSCs, the shift observed in recurrent GBM tissues is likely due to the reported contribution of the immune microenvironment to mesenchymal signatures (Hoogstrate *et al*., 2023; Wang *et al*., 2017). Nevertheless, upregulation of subsets of makers associated with mesenchymal or neuronal phenotypes was evident in GSCs acquiring secondary TMZ or IR resistance. Indeed, changes emerging in therapy-surviving GSCs can be subtle and difficult to detect by global transcriptomics, yet still have a powerful impact on GSC biology.

This is the case of RTKs, which were often more highly expressed in GSCs emerging from therapies, particularly after IR and in cases of secondary resistance. These findings align with previous mechanistic studies showing the prominent role of EGFR and MET in supporting GBM therapeutic resistance (De Bacco *et al*., 2016; Squatrito & Holland, 2011). Notably, in our dataset the increase in RTK protein was not accompanied by increased mRNA expression, suggesting sustained activation of post-transcriptional mechanisms, and it was reflected functionally by increased responsiveness to the corresponding growth factors essential for propagation. This observation highlights that the ability of GSCs to survive genotoxic treatments likely depends not only on mechanisms directly addressing DNA damage and repair (e.g., MGMT activity or MMR inactivation), but also on survival pathways orchestrated by RTK upon stimulation by growth factors present in the microenvironment. The RTK pathway and its intertwining with apoptotic control may represent a strength of recurrent GSCs and a potential weakness, if targeted appropriately in combination therapies.

Overall, res-GSC families provide a within-patient, therapy-defined system to separate TMZ- from IR-driven resistance trajectories and capture (i) durable genetic escape under TMZ (MMR loss), (ii) MGMT-independent drug tolerance, (iii) therapy-associated chromosomal evolution, and (iv) convergent adaptive programs such as RTK remodeling. By enabling controlled, pressure-specific mechanistic testing, this framework complements primary–recurrent GBM tissue profiling and helps identify vulnerabilities to delay, prevent and treat GBM recurrence.

## METHODS

### Patient samples

For derivation of Res-GSC families, glioblastomas (GBMs) were obtained from patients that underwent surgery at Città della Salute e della Scienza (University of Torino, Italy) and were processed at the Candiolo Cancer Institute between January 2014 and March 2020. Other GSC cultures (neurospheres) used for set-up experiments (Fig. EV1) were derived from GBMs that underwent surgery and were processed at the Fondazione IRCCS Istituto Neurologico C. Besta (Milan, Italy) (BT302: April 2009, BT483: February 2012, and BT205: August 2008) and were previously described (De Bacco *et al*, 2012; De Bacco *et al*., 2016). All GSCs were regularly tested and authenticated through STR analysis with PowerPlex 16 HS System (Promega) performed by the research group at the Candiolo Cancer Institute. All patients were recruited under protocols approved by the institutional Ethical Committees at Città della Salute e della Scienza (University of Torino, Italy) or at the Fondazione IRCCS Istituto Neurologico C. Besta (Milan, Italy). All patients gave their informed written consent, and the studies were carried out following the Declaration of Helsinki. Before processing, all patient data and samples were de-identified and are reported in Table EV1 and in a previous work (De Bacco *et al*., 2023b). For the genetic comparisons with TCGA GBM cohort, data for were obtained from the public cBioPortal website (Glioblastoma Multiforme TCGA Firehose Legacy, July 2022, https://www.cbioportal.org/) (Cerami *et al*, 2012).

### Res-GSC derivation

Res-GSC family members were derived starting from primary GBMs, surgically removed by aspiration, and processed to single cell suspension according to a previously published protocol (De Bacco *et al*., 2023a). Briefly, aspirator’s content was collected in 50 mL conical tubes, centrifuged RT 10 min at 270 g, and red blood cells were lysed by treatment with ACK for 15 min. The pellet was incubated with collagenase type 1 (Thermo Fisher) at 37°C for 20 min, mechanically dissociated by up and down pipetting with a sterile 18G needle in a 1 mL syringe and subjected to sequential 100 mm and 70 mm filtrations. Cellular suspension was equally distributed in three 75 cm^2^ flasks in Dulbecco’s modified Eagle’s medium/F-12 (Thermo Fisher) with the addition of 2 mM glutamine (Sigma), antibiotic and mycotic solution (1:100, Sigma), B-27 plus (1:50, Thermo Fisher), human recombinant epidermal growth factor (EGF), basic fibroblast growth factor (FGF2), hepatocyte growth factor (HGF) and platelet-derived growth factor (PDGFBB) at 20 ng/ml each (Peprotech). After a short period in culture allowing cell recovery, one flask was treated with temozolomide (TMZ, Sigma) at 50 μM every three days for three times, another flask was irradiated at 2 Gy dose for five consecutive days by using a 200-kV X-ray blood irradiator (Gilardoni, Lecco, Italy), with a 1 Gy/ minute dose rate; the third flask remained untreated. Flasks were maintained in normoxic condition (20% O_2_, 5% CO_2_) at 37°C. A simplified scheme of derivation procedure is provided in Fig. 1A. Part of the same cellular suspension was collected for DNA analysis. GSCs were propagated in derivation medium, were considered derived when reaching passage 5, and fully stabilized after 10 passages.

### Nucleic acid extraction

Genomic DNA (gDNA) was extracted from whole GBM cell suspensions and GSC cultures using the ReliaPrep™ gDNA Tissue Miniprep System (Promega) according to manufacturer’s protocol. Total RNA for RNA sequencing was extracted from GSC cultures using MaxwellRSC symply RNA Cells kit, and for qPCR with Maxwell® RSC miRNA tissue kit (Promega) according to manufacturer’s instructions. Concentrations and purity of nucleic acid were assessed spectrophotometrically using A260/A280 absorbance ratios with Nanodrop De Novix Ds-11 spectrophotomer (Resnova). When undergoing Next Generation Sequencing, nucleic acid concentration was also evaluated using the proper Qubit RNA BR (or HS) or dsDNA BR (or HS) Assay kit (Thermo Fisher).

### Droplet digital PCR (ddPCR)

To estimate tumor purity in original cell resuspension, *TERT* analysis was performed according to Corless et al.(Corless *et al*, 2019). Tumor cell purity was calculated multiplying *TERT* variant allele frequency (VAF) *2. To detect possibly rare gene alterations in original UA, ddPCR was performed using probe–based assays with ddPCR Supermix for Probes (no dUTP) (Biorad), according to manufacturer’s instructions. In all cases, droplets were generated using AutoDG Droplet Digital PCR System (Biorad) and analyzed on a QX200 Droplet Digital PCR System using QX Manager Software Standard Edition, Version 1.2 (Biorad). For ddPCR probes see Dataset EV2.

### NGS target panel analysis

Library preparation was performed starting from 400 ng of DNA, firstly fragmented by using the M220 Focused-ultrasonicator (Covaris), followed by a clean-up step with an optimized ratio volume of AMPure XP beads (Beckman Coulter). Subsequent end-repair step and dA-tailing reaction of blunt-ended DNA fragments was performed by means of NxSeq AmpFREE Low DNA Library Kit (Lucigen) with small adjustments to increase the efficiency of the reactions. Adaptor ligation step has been performed with the same Lucigen’s kit, by using xGen Stubby Adapter (IDT) as adaptors. After clean-up step with AMPureXP beads (Beckman Coulter), samples have been amplified (KAPA HiFi HotStart ReadyMix PCR Kit, Roche) concomitantly introducing unique sample barcodes (xGen Stubby Adapter-UDI Primers, IDT). Before target enrichment, QC of post-PCR libraries were checked by means of Qubit dsDNA BR Assay kit (Thermo Fisher) and of 2100 Bioanalyzer with a High-Sensitivity DNA assay kit (Agilent Technologies). The target of interest for GBM-custom panel design has been defined starting from the identification of genes relevant for tumorigenesis, evolution and emergence of drug resistance in GBM, thus including all coding regions of 75 genes (De Bacco *et al*., 2023b). Equal amounts of post-PCR libraries (750 ng) were pooled, for a maximum of 8 samples per pool, and subjected to the GBM-panel target enrichment with xGen Hybridization and Wash Kit (IDT plus xGen Universal Blockers - TS Mix (IDT)) following manufacturer instruction, except for the choice to perform over-night hybridization to increase the on-target capture. A further amplification of the libraries has been performed with KAPA HiFi HotStart ReadyMix PCR Kit (Roche) and xGen Library Amplification Primer Mix (IDT), thus reaching the needed number of final libraries. Final libraries were quantified by means of Qubit dsDNA HS Assay Kit (Thermo Fisher) and their fragment distribution evaluated using High-Sensitivity DNA assay kit (Agilent Technologies). Equal molar amounts of DNA libraries were pooled and, based on the total number of samples ready to be analyzed, sequenced using Illumina MiSeq or NextSeq500 sequencer (Illumina).

Data analysis was performed using a bioinformatic pipeline previously described (Corti *et al*, 2019; Crisafulli *et al*, 2019). A metanormal was built from fastQ files obtained by 10 PBMC samples sequenced using the same laboratory procedures (Battuello *et al*, 2024; Crisafulli *et al*., 2022; Pignochino *et al*, 2021). Alignments from metanormal and whole GBM cell population samples were compared to identify mutations/indels in tumor and metanormal samples. Somatic alterations were present only in tumor while germline ones were common to both samples. NGS artifacts were further filtered following the methods previously described (Crisafulli *et al*., 2019; Crisafulli *et al*., 2022). Then only variants with 5% significance level obtained with a Fisher’s exact test, supported by a minimum of 4 mutated reads in regions with 53 minimum depth and with allele frequency >2.5% were considered. Indels were called using Pindel tool in both alignments and only somatic indels with fractional abundance >10% were reported according to methodology previously reported (Crisafulli *et al*., 2019; Crisafulli *et al*., 2022).

Gene CN variations analysis was performed in the matched samples (Tumor vs. Metanormal) for each patient as previously reported (Crisafulli *et al*., 2019; Crisafulli *et al*., 2022). Pathogenicity of all identified variants was manually curated curated using MutationTaster2021(Steinhaus *et al*, 2021), Alphamissense (Tordai *et al*, 2024) and the Catalog Of Somatic Mutations In Cancer(Sondka *et al*, 2024). When appropriate alterations were validated by qPCR, or Sanger Sequencing and/or droplet digital PCR (ddPCR).

### Gene copy number evaluation

Gene copy number (CN) analysis was assessed by real-time PCR, using TaqMan Universal PCR Master MIX and the ABI PRISM 7900HT sequence detection system (Thermo Fisher). Primers and probes for TaqMan CN assays are reported in Dataset EV2. Relative gene CN data were calculated by normalizing against endogenous controls (*RNaseP* and/or *APOA1* and/or *GREB1*). Normal diploid human gDNA (from PBMCs) was used as calibrator to obtain the ΔΔCt. The CN of each gene was calculated with the formula 2×2^-ΔΔCt^. To discriminate between real *EGFR* amplification and chr7 polysomy, the calculated CN was normalized vs. CN of a usually not amplified reference gene mapped on chr7 (*HGF*).

Genes alterations were defined as follow: amplification was defined when CN was > 5 (for *EGFR* amplification was defined when CN was > 3 + *HGF* CN); CN gain was defined for CN among 3 and 5; heterozygous deletion was defined when CN is < 1.5; homozygous deletion was defined when CN < 1 in tumor tissues, while corresponded to the absence of target gene PCR product in the presence of control gene PCR product in growing cells.

### Sanger sequencing

DNA was amplified using Platinum Taq DNA Polymerase (Thermo Fisher) and specific primer pairs (Dataset EV2). PCR conditions were as follows: 95°C for 3’; 3 * [95°C for 15″, 64°C for 30″, 70°C for 1’]; 3 * [95°C for 15″, 61°C for 30″, 70°C for 1’]; 3 * [95°C for 15″, 58°C for 30″, 70°C for 1’]; 37 * [95°C for 15″, 57°C for 30″, 70°C for 1’]; and 70°C for 5’. PCR products have been checked using 1.5% agarose gel and purified using illustra™ ExoProStar 1-Step (Merk) following manufacturer indications.

Cycle sequencing was performed using BigDye Terminator v3.1 Cycle Sequencing kit (Thermo Fisher). Sequencing products were purified using CleanSEQ Dye-Terminator Removal Kit (Beckman Coulter) and analyzed with a 3730xl Genetic Analyzer (Thermo Fisher). Data were visualized by Chromas Lite 2.6.6 software and compared with reference sequences from the Homo sapiens assembly GRCh37.

### EGFRvIII analysis

mRNA from whole GBM cell populations have been analyzed for the expression of EGFRvIII variant. mRNA was retro-transcribed using high-capacity cDNA reverse transcription kit (Thermo Fisher) following manufacturer’s instruction. Three primer pairs were designed: 1 specific for wild type EGFR, to be used as reaction control, and the other two for EGFRvIII (see Dataset EV2) to be used in three PCR reactions. cDNA was amplified using Platinum Taq DNA Polymerase (Thermo Fisher). PCR conditions were as follows: 93°C for 3’; 40 * [93°C for 15’’, 60°C for 30’’], 72°C for 30’’; final extension 72°C for 3’. PCR products were evaluated on a 2% agarose gel.

### *MGMT* promoter methylation analysis

The evaluation of *MGMT* promoter methylation in tumor diagnostic sample, (formalin-fixed paraffin-embedded samples, FFPE) was performed by pyrosequencing assay. Approximately 200-500 ng total DNA was subjected to bisulfite conversion and pyrosequencing analysis using the MGMT plus kit (Diatech Pharmacogenetics, Italy) and the PyroMark Q96 ID system (Qiagen, Valencia, CA, USA). The pyrosequencing assay was performed as described earlier (Shaw *et al*, 2006). The primers used for amplification of bisulphite-treated DNA were forward: 5′-GGGATAGTTGGGATAGTT-3′ (the first g avoids formation of hairpin loops) and reverse: 5′-biotin-ATTTGGTGAGTGTTTGGG-3′ giving a 99-bp amplicon (see also Dataset EV2). The PCR analysis was performed in duplicate in 25 μl reaction volume, containing 300 pmol each forward and reverse primer, 2 μl 10 × buffer, 160 μM dNTPs, 0.5 U HotStar Taq polymerase (Qiagen) and 1–2 μl bisulfite-treated DNA. The PCR conditions were as follows: 95°C for 15’; 40 * [94°C for 30’’, 50°C for 45’’, 72°C for 30’’] and 72°C for 10’. 3 μl of PCR products were analyzed on a 2% agarose gel, the remaining 22 μl was subjected to pyrosequencing on a PSQ96MA System (Biotage, Uppsala, Sweden) using the primer 5′-GGATATGTTGGGATAGT-3′and PyroGold reagents (Biotage). The Pyro Q-CpG software 1.0.9 (Biotage) was used to analyze data.

Pyrosequencing yields data for 10 CpG sites within the *MGMT* promoter. For data analysis, the percentage methylation obtained for each CpG was averaged across the 10 CpGs in duplicate PCR reactions (average methylation per sample). For comparisons with clinical data, glioblastomas were considered methylated if they had at least one sample with average methylation ⩾9% (⩾mean+2 s.d. for non-neoplastic brain) in more than one independent bisulphite modification. The average methylation per case was calculated by averaging the average methylation per sample for methylated samples for that case. According to Dunn et al. (Dunn *et al*, 2009), GBMs were considered methylated if they had an average methylation ≥ 9.

The evaluation of *MGMT* promoter methylation in whole GBM cell suspensions and established GSC cultures was performed using ddPCR after DNA treatment with bisulfite. For our analysis, bisulfite conversion was done using the EZ DNA Methylation Gold kit (Zymo Research) according to manufacturer protocol. For ddPCR reaction 5-10µ of DNA template was added to 10 µL of ddPCR Supermix for Probes (Bio-Rad) and 5 µL of the primer and probe mix (the fluorophores used are FAM and HEX). Droplets were generated using AutoDG Droplet Digital PCR System (Biorad) and analyzed on a QX200 Droplet Digital PCR System (Bio-Rad). Samples were considered methylated if the value of methylation was ≥30%. For whole GBM cell populations, normalization on the percentage of tumor cells (based on *TERT* promoter mutation VAF, see above) was done using the formula: *MGMT* methylation % on tumor = (100 * observed *MGMT* methylation %) / tumor cell %.

### Chromosome G banding

Cells were synchronized 48-72h after plating using out by Synchroset (Euroclone) according to manufacturer’s instructions; mitotic spindle was blocked using Colcemid Solution (10 μl/ml) (Euroclone) for 1 h. Chromosome harvesting was carried out according to standard procedures. Briefly, cells were incubated in hypotonic solution (KCl 0,075 M) for 10 min and fixed in methanol-glacial acetic acid (3:1). Cell suspensions were dropped on slides and dried using specific conditions for optimal chromosome spreading. G banding was achieved by incubating slides in 2×SSC (Merk) at 65 °C for 2 min and then stained with Wright’s solution for 2 min (Merk).

Acquisition of metaphases was performed using an Olympus BX61 microscope (Olympus Corporation) and the analysis was carried out by CytoVision software 7.2 (Leica Biosystems). A mid-range resolution of 300 bands was achieved. The anomalies were reported according to the International System for Human Cytogenomic Nomenclature (2024).

### Chromosome M-FISH

Chromosome analysis by Multicolor-Fluorescence In Situ Hybridization (M-FISH) was performed on GSCs through using 24XCyte Human Multicolor FISH Probe (MetaSystems Probes). Probes and metaphase chromosomes denaturation and hybridization were performed according to manufacturer’s instructions. Briefly: slides were incubated at 70 °C in saline solution (2× SSC), denatured in 0.07N NaOH, dehydrated in an ethanol series and air-dried. Probe denaturation was carried out by incubating probe cocktail at 75°C (±1°C) for 5min, 2 minutes on ice and at 37°C (±1°C) for 30 min. 10 μl of the denaturated probe cocktail were put onto slides and hybridization for 24h at 37 °C followed. Subsequently, slides were washed with post-hybridization buffers and counterstained with 10 μl of DAPI/antifade (MetaSystems Probes). The Metafer System and the Metasystems ISIS software (MetaSystems Probes) were used for signal detection and metaphase analysis. At least ten metaphases of each culture were studied for each cell line.

Chromosomal aberration acquired by TMZ-GSCs or IR-GSCs were counted considering the prevailing clone for each GSC family member as follows: a score of 1 has been assigned for each type of structural aberration acquired or lost with respect to CTRL-GSC (if multiple chromosomes display the same alteration, the assigned score remains 1).

### Evaluation of MGMT by FISH

FISH analysis was performed to verify the proper chromosome location of MGMT (10q26.3) gene. Clones RPCI11-333H4 (proximal to MGMT) and RPCI11-300B2 have been provided as a courtesy by Prof. Mariano Rocchi (University of Bari) and expanded in vitro in liquid broth supplemented with 12.5 ug/ml Cloramphenicol. BAC DNA was prepared according to Rondon et al.(Rondon *et al*, 1999).

1µg of each BAC DNA has been directly labelled using the Nick Translation Reagent Kit using 2.5μl of 0.2mM SpectrumRed-dUTP or 3.5μL of 0.2mM SpectrumGreen-dUTP (Abbott Molecular Inc., Abbott Park, IL) following manufacturer instruction, precipitated and resuspended in 50µL of TE buffer. 10µL of each probe was pooled, precipitated using 10µg of carrier DNA (sheared ssDNA, Thermo Fisher) and 6 µg of Human COT DNA (Roche), and resuspended in 150μl Hybridization solution (2xSSC, 50% formamide, 20% dextran sulfate). Slides and probes were then processed as described for M-FISH.

### MSI status evaluation

MSI status has been evaluated using the OncoMate® MSI Dx Analysis System (Promega) following manufacturer indications. PCR products have been analyzed with a 3730xl Genetic Analyzer (Thermo Fisher).

### Gene expression analysis by quantitative real-time PCR (qPCR)

Purified mRNAs were reverse transcribed starting from 150 or 250 ng of total RNA and using High-Capacity cDNA Reverse Transcription Kit (Thermo Fisher). Real-time PCR for evaluation of *RAD51* gene mRNA expression was performed using primer and probe sets (Thermo Fisher) listed in Dataset EV2, with TaqMan Universal PCR Master Mix and an ABI PRISM 7900HT sequence detection system (Thermo Fisher). Expression levels were normalized against endogenous control (β2 microglobulin). Control (not irradiated cells) cells were used as calibrators. Expression levels were reported as relative expression. Results are shown as mean ± of SEM of three independent experiments.

### Bulk gene expression profiling and Single-Sample Gene Set Enrichment Analysis (ssGSEA)

Poly(A)-enriched mRNA sequencing was performed by Novogene following standard procedures for Human mRNAseq. Raw gene-level count data were generated processing fastq files using the function rnaseqCounts implemented in docker4seq(Beccuti *et al*, 2018) using the GRCh38.p7 reference assembly. All analyses were conducted in R version 4.4.2 (R Core Team, 2024) using the packages readxl version 1.4.5, data.table version 1.18.0, and AnnotationDbi version 1.68.0. Gene identifiers were annotated using the org.Hs.eg.db version 3.20.0. Custom ssGSEA scripts were sourced locally. All computational steps were executed through documented R scripts to support reproducibility. Transcriptional subtype-specific signatures were defined from previously curated programs: Wang et al.(Wang *et al*., 2017) (Classical, Proneural, Mesenchymal); Garofano et al.(Garofano *et al*., 2021) (GPM, MTC, NEU, and PPR); and Neftel et al.(Neftel *et al*., 2019) (MES1, MES2, NPC1, NPC2, OPC, AC). Gene sets were provided as Excel tables and as .mod files compatible with ssGSEA. For each signature, overlap between the curated gene list and the processed expression matrix was evaluated prior to scoring. Sample classification was performed using a custom ssGSEA implementation based on the method described by Wang et al.(Wang *et al*., 2017), including equivalent distribution resampling. The normalized expression matrix and subtype-specific .mod files were used as input. Enrichment scores were computed independently for each sample (n=27) using 10000 permutations to assess statistical robustness. ssGSEA was implemented using custom R scripts (msig.library.12.R and runSsGSEAwithPermutationR3.R), adapted from Wang et al. (Wang *et al*., 2017)

### γH2AX immunofluorescence assay

GSCs were mechanically dissociated at single cell level and about 10^5^ cells were seeded in 6-well plate. 6 days after seeding, cells were irradiated with a single 5 Gy dose and were fixed 2, 3 and 4 hours after with PAF 4% for 20 min. Fixed cells were washed with milliQ water, resuspended in 100 µL of water and seeded on poly-lysinated slides. Cells were then washed with PBS 3 times for 5 min, permeabilized with HEPES-Triton for 5 min, washed with PBS for 5 min and saturated with PBS-BSA 5% for 1 h. Primary antibody γH2AX (Cell Signaling Technology, cat#9718), diluted 1:200 in PBS-BSA 0.5%, was incubated for 2 h RT. Appropriate AlexaFluor conjugated secondary antibody was used (Thermo Fisher, 1:1000 in PBA-BSA 0.5%) for 1 h RT. DAPI was added for 30 sec. Images were captured using LASV4.2 software on a LEICA SPEII confocal microscope. The percentage of γH2AX positive cells has been measured by counting positive cells/nuclei (DAPI pos) in at least 10 HMFs.

### Western blotting

GSC proteins were extracted with boiling Laemli Buffer or RIPA buffer. Total amount of proteins obtained was quantified using the BCA System (Pierce). 10 to 15 ug of proteins were loaded on a 4-15/4-20% pre-casted polyacrylamide gel (Biorad) and transferred on nitrocellulose. Commercial and home-made (anti-MET (Prat *et al*, 1998)) primary antibodies used were as follows: anti-MGMT (Cell Signaling Technology, cat#2739), anti-RAD51 (Cell Signaling Technology, cat#8875), anti-MRE11 (Cell Signaling Technology, cat#4895), anti-RAD50 (Cell Signaling Technology, cat#3427), anti-NBS1 (Cell Signaling Technology, cat#14956), anti-CD44 (Cell Signaling Technology, cat#5640), anti-Vimentin (Millipore, cat#CBL202), anti-CHI3L1/YL40 (Cell Signaling Technology, cat#47066), anti-STAT3 (R&D, cat#MAB1799), anti-YAP/TAZ (Cell Signaling Technology, cat#8418), anti-OLIG2 (Millipore, cat#AB9610), anti-SOX10 (Santa Cruz Biotechnology, cat#sc-365692), anti-GFAP (Santa Cruz Biotechnology, cat# sc-6171), anti- S100β (Invitrogen, cat#PA5-78161), anti-Doublecortin (Abcam, cat#ab207175), anti-NeuroD1 antibody (Abcam, cat#ab213725), anti-β3-Tubulin (Cell Signaling Technology, cat#4466), anti-NeuN/RBFOX3 (Invitrogen, cat#702022), anti-EGFR (Cell Signaling Technology, cat#4267), anti-PDFGFRα (Cell Signaling Technology, cat#3174), anti-PDGFRβ (Cell Signaling Technology, cat#4564), anti-Phospho-Chk2 (Cell Signaling Technology, cat#2661), anti-CHK2 (Cell Signaling Technology, cat#3440), anti-p21 (Millipore, cat#05-655). Antibodies were visualized with appropriate horseradish peroxidase-conjugated secondary antibodies (Jackson Lab) and enhanced chemiluminescence system (Promega). Blot images were captured using the ChemiDoc TouchTM Imaging System (Biorad) with Image Lab software (Biorad). H3 (Millipore, cat# 05-499), β-actin (Cell Signaling Technology, cat#8457), vinculin (Sigma-Aldrich, cat#V9131) and calnexin (Cell Signaling Technology, cat#2679) were used as protein loading control where indicated.

### Flow-cytometric analyses

All cases underwent extensive immunophenotyping via multiparametric flow cytometry. From growing GSC cultures, single cell suspension was obtained by mechanical disaggregation, immediately counted and evaluated for cell viability with Biorad TC20 automated cell counter, in order to verify that viability was more than 70% and that total number of viable cells was greater than 1.5×10^6^. Cells were then washed with PBS/BSA 1% and resuspended in the same solution at 2×10^5^ cells/100 µL, to prevent nonspecific binding of detection antibodies to Fc receptors (FcR) on immune cells, ensuring antigen-specific detection and minimizing false-positive signals. 100 µL of samples were directly added to a monoclonal antibody mixture, incubated 20 minutes RT, washed twice with PBS/BSA 1%, resuspended in 300 μL of PBS/DAPI 0.2X and immediately acquired. For evaluation of surface expression, the immunophenotypic sample acquisitions were performed by a CyAn ADP (Beckman Coulter) equipped with 3 solid state lasers (488 nm, 405 nm and 635 nm) and 9 colors. Analyses were performed by Summit 4.3 software (Beckman Coulter). For evaluation of intracellular expression, the immunophenotypic sample acquisitions were performed by a Cytoflex (Beckman Coulter) equipped with 3 solid state lasers (488 nm, 405 nm and 638 nm) and 8 colors. Analyses were performed by CytExpert 2.6 software (Beckman Coulter). The antibodies used for immunophenotype characterization were as follows: anti-EGFR (BioLegend, cat#352908), anti-HGFR/c-MET (BioTechne, cat#MAB3582), anti-PDFGFRα (BioLegend, cat#323506), anti-PDGFRβ (BioLegend, cat#323608), anti-FGFR1 (Immunological Sciences, cat#AB-844949), anti-FGFR2 (BioTechne, cat#FAB684A), anti-FGFR3 (BioTechne, cat#FAB6852A), anti-FGFR4 (BioTechne, cat#FAB766A). Death cells were excluded from analysis using DAPI (Roche) staining.

FC score for RTK surface expression was calculated as the sum of the score for the percentage of positive cells and the mean fluorescent intensity (MFI). Specifically: a score of 0 was assigned if the % of positive cells ≤5, or MFI=0; a score of 1 if the % of positive cells >5 and ≤ 20, or MFI<25; a score of 2 if the % of positive cells >20 and ≤ 40, or MFI<100; a score of 3 if the % of positive cells >40 and ≤ 60, or MFI<250; a score of 4 if % of positive cells >60 and ≤ 80, or MFI<500; a score of 5 if the % of positive cells >80 and ≤ 100, or MFI≥500.

### Cell cycle evaluation (pospho-H3 staining)

To thoroughly investigate the distribution of cells across different phases of the cell cycle and any variations induced by treatment, flow cytometric analysis was employed. A total of 1×10⁶ cells were fixed on ice in a solution of 10% methanol and 37% paraformaldehyde, followed by staining in a permeabilization buffer containing 5% saponin, 100 mM HEPES (pH 7.4), 1.4 M NaCl and 25 mM CaCl₂. To discriminate and quantify cells between the G2 and M phases of the cell cycle, the phosphorylation of histone H3 at Ser10 was detected, as this correlates with the G2 to M phase transition(Juan *et al*, 1998; Prigent & Dimitrov, 2003). The cells were stained with an anti-phospho H3 (Ser10) antibody conjugated with AlexaFluor® 488 (Sigma Aldrich, cat#FCMAB104A4). Finally, staining with 1 µl of FxCycle Violet (Thermo Fisher Scientific) in 500 µl of permeabilization buffer enabled precise quantification of DNA content for cell cycle analysis by exploiting stoichiometric binding to DNA (Darzynkiewicz *et al*, 2004).

### Cell viability assays

Approximately 10^3^ cells per well were plated in a 96-well plate 24 hours before starting the treatment, and viability was measured by CellTiter-Glo Luminescent Cell Viability Assay (Promega) using a GloMax 96 Microplate Luminometer (Promega) as detailed below.

To evaluate TMZ doses able to discriminate between sensitive and resistant cells and, subsequently, cell response to chemotherapy, cells were treated at day 0, 3 and 6 with TMZ (Sigma), with doses ranging from 1 to 50 µM, and cell viability was measured 0, 3, 6 and 9 days after the beginning of the treatment (cell viability at day 9 was shown).

To evaluate cell response to IR, cells were irradiated with a dose of 2 Gy administered for three consecutive days or with a single dose of 5 Gy. Cell viability was measured 48 hours after the last treatment. In both cases, data are reported as mean ± SEM of at least two independent experiments.

### Clonogenic assay under therapeutic pressure

Clonogenic assay was performed as described earlier(Franken *et al*., 2006). Cells were seeded at the concentration of 5-25 cells per well in 96-well plates in the culture medium. To evaluate cell response to chemotherapy, TMZ stimuli at 5 and 50 μM were administered once 24 hours after seeding. To evaluate cell response to IR, cells were irradiated with a dose of 2 Gy administered once (2 Gy) or for three consecutive days (2 Gy × 3) or with a single dose of 5 Gy. Neurospheres or foci generated in all conditions were counted when vehicle/mock-treated cells displayed formed neurospheres/foci. Residual clonogenic potential after treatment (i.e. surviving fraction) was calculated normalizing the number of neurospheres/foci in each condition with the number of neurospheres/foci in vehicle/mock-treated wells and considering the plating efficiency. Data are reported as mean ± SEM of at least two independent experiments.

### Extreme limiting dilution assay (LDA)

Cells were seeded at the concentration of 100, 50, 25, 10, 5 and 1 cell per well (10 replicates for each condition) in 96-well plates in the culture medium. Wells containing neurospheres/colonies were counted and data were analyzed with ELDA software (http://bioinf.wehi.edu.au/software/elda/) (Hu & Smyth, 2009). For p1 passage, neurospheres were collected and dissociated, and derived cells were seeded at the same concentrations described above. Data reported have been obtained from passage p1 and are plotted as mean ± SEM of at least two independent experiments.

### Cell proliferation assay

Cell proliferation was measured by CellTiter-Glo Luminescent Cell Viability Assay (Promega) using a GloMax 96 Microplate Luminometer (Promega) at day 0 (ctrl), and at the indicated time points after treatment. To evaluate the responsiveness to single GF and the ability to growth in the absence of exogenous stimuli, 10^3^ dissociated cells per well were seeded in a 96-well plate at day -1 in a medium devoid of any GF; on day 0 cells were stimulated with the indicated GF. Data were reported as mean ± SEM of at least two independent experiments. Growth responsiveness (GF resp) score was calculated assigning a value of 1 to each growth factor inducing a cell proliferation greater than 1.5 and summing up all the growth factors’ contributions.

### Fluorescence-based multiplex flow-cytometric host cell reactivation assay (FM-HCR)

Plasmids for MMR activity evaluation through FM-HCR (Nagel *et al*., 2014; Piett *et al*., 2021), included pMax_BFP and pMax_mOrange control plasmids, and mOrange_GG, which is a modified pMax_mOrange plasmid that has a site-specific G:G mismatch. Repair of the mismatch results in a change of sequence in the transcribed strand and leads to expression of mOrange fluorescent protein. As expected, mOrange_GG yielded a robust fluorescent signal only when MMR proficient cells were transfected. Two parallel co-tranfections were carried using 2 wells of a 12 well plate, with 1.25×10^5^ cells were plated in 1 mL of complete medium without antibiotic antimitotic solution, implemented with 5% of fetal bovine serum (Euroclone) each. 24 h after plating, cells were transfected using, for each well, 100 µL of Opti-Mem containing 850 ng of carrier DNA (sheared ssDNA, Thermo Fisher) and 4 µL of TransIT-X2 Dynamic Delivery System (Mirus Bio) and plasmid DNA: 25 ng of mOrange_GG + 25 ng of pMax_BFP (co-transfection 1) or 25 ng of pMax_mOrange + 25 ng of pMax_BFP (co-transfection 2). 24 h post transfection, cells were harvested by trypsinization, washed in 1X PBS, and each sample was resuspended in 250μL of a solution containing LIVE/DEAD Fixable Near-IR Dead Cell (ThermoFisher) diluted 1:1000 in PBS/BSA 1%. Cells were incubated for 30 min at room temperature in the dark on a seesaw rocker. After washing in 1X PBS to remove excess dye, samples were resuspended in 200 μL of 1X PBS and immediately analyzed by FC. Sample acquisition was performed using a CytoFLEX LX (Beckman Coulter) equipped with 5 solid-state lasers (375 nm, 405 nm, 488 nm, 561 nm, 638 nm). Analyses were performed using CytExpert software (Beckman Coulter). MMR efficiency (Z, percent reporter expression) was calculated as follow: Z=X/Y*100, where X= (mOrange_GG normalized count * mean mOrange_GG intensity) / (BFP normalized count * Mean BFP intensity) and Y= (mOrange normalized counts * mean mOrange intensity) / (BFP normalized counts * Mean BFP intensity). In each case, mean intensity was for the cells that were positive for BFP or mOrange, respectively. BFP reporter expression provides a control for transfection efficiency. Percent reporter expression is an estimate of the percentage of mismatch-containing mOrange_GG plasmids that have been repaired.

## QUANTIFICATION AND STATISTICAL ANALYSIS

Where indicated, data were expressed as mean ± standard error of the mean (SEM) of at least two independent experiments. p<0.05 was considered statistically significant. Statistical comparisons were performed using the parametric Student t test, Wilcoxon matched-pairs signed rank test, Mann-Whitney test, Fisher’s exact test and One Sample Wilcoxon test as reported. Normality of the data was assed using Shapiro-Wilk Test. All statistical tests were performed using Prism v8.0 software (GraphPad). Where indicated, ELDA software (http://bioinf.wehi.edu.au/software/elda/) was used.

## MATERIALS AND DATA AVAILABILITY

### Materials availability

GSCs are available from the corresponding author upon completed material transfer agreement after request by qualified academic investigators for non-commercial purposes.

### Data availability

Human Next Generation Sequencing data (DNA sequencing and bulk RNA sequencing) will be available at the European Nucleotide Archive (ENA) of European Bioinformatics Institute (EBI) under project accession number PRJEB107184, study ERP188209 (available at https://www.ebi.ac.uk/ena/browser/search) and will be made public upon acceptance for publication.

RNAseq data are also reported as TPMs (Dataset EV1). Other source data will be deposited at Mendeley and will be made public upon acceptance for publication. Any additional information required to reanalyze the data reported in this paper is available upon request from the corresponding author.

## AUTHOR CONTRIBUTIONS

Conceptualization, M. Prelli., F.D.B., F.O. and C.B.; methodology, M. Prelli, F.D.B. and F.O.; formal analysis, M.Prelli, F.D.B., E.C., S. Mahmoudi, R.C. M. Panero, L.C., G.C. and F.O.; investigation, M. Prelli, F.D.B., E.C., S. Maniscalco, G.B., G.R., M. Panero, E.B., A.B. and M.M.; resources, Z.D.N., L.B., P.C., P.Z., F.C. and D.G.; writing–original draft, M. Prelli, F.O. and C.B.; writing–review & editing, M. Prelli, F.D.B., F.O., G.C. and C.B.; supervision and funding acquisition, C.B.

## ACKNOWLEDGEMENTS

We thank A. Melcarne for neurosurgeries, D. Cantarella, S. Gilardi, B. Martinoglio, R. Porporato, N. Calandra for technical help and D. Gramaglia and M. Del Boccio for assistance. This study has received funding from AIRC under IG 2023 – ID. 28836 – P.I. Boccaccio Carla, Italian Ministry of Health RC 2023-2025 and Comitato per Albi98 (to C.B.), and American Cancer Society Grant RSG-22-038-01-DMC (to Z.D.N.).

---

**Fig. EV1.**
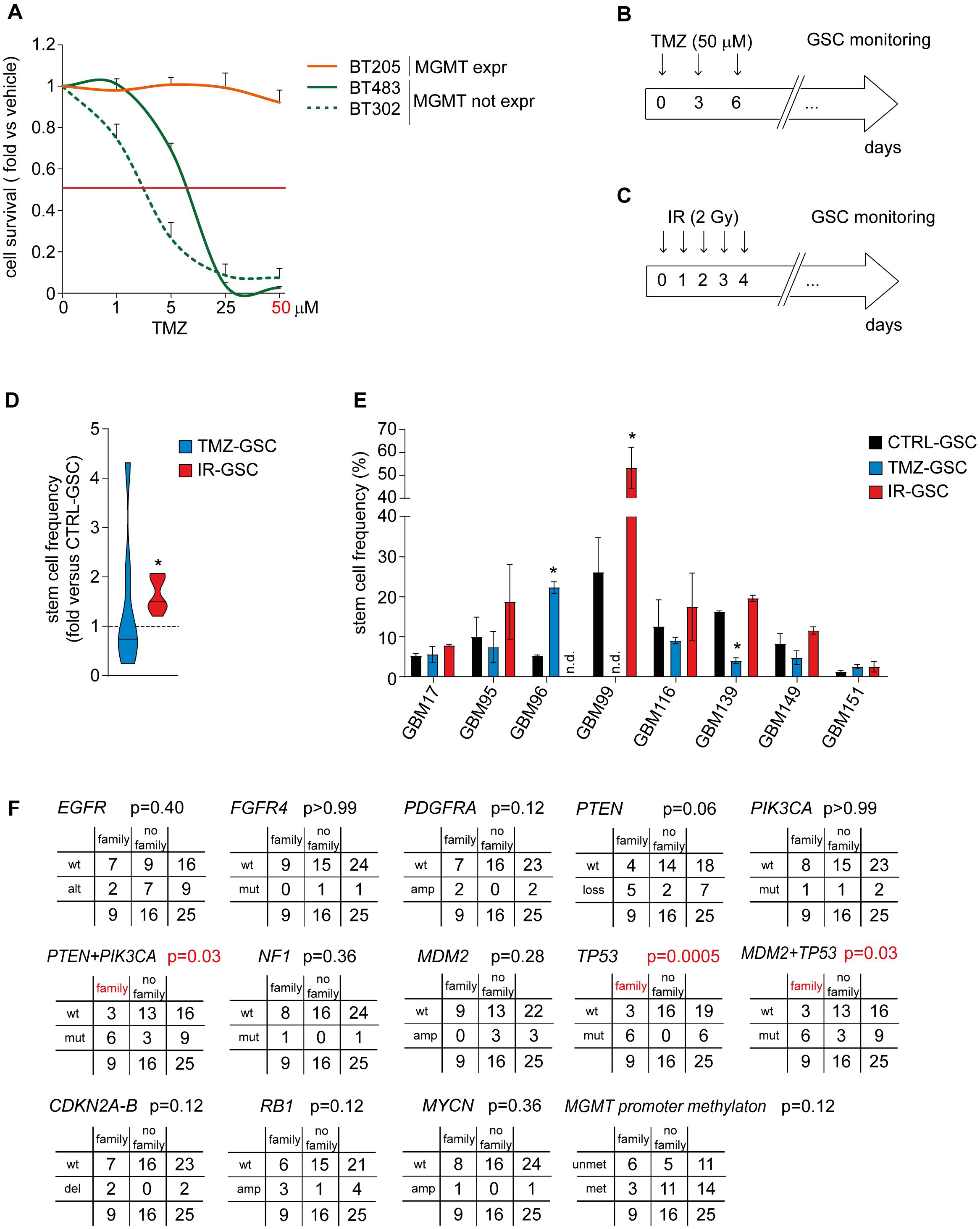
Protocols for res-GSC family derivation under therapeutic pressure and correlations with original GBM genetic features. (**A**) Dose-response to temozolomide (TMZ) measured as cell survival (fold vs. vehicle) in two representative primary sensitive GSCs (BT302 and BT483, MGMT-neg, not expressing the dealkylating enzyme methylguanine methyltransferase) and a primary resistant GSCs (BT205, MGMT-pos, expressing MGMT). Red line indicates 50% cell viability vs. vehicle. Vehicle: cells treated with DMSO. n = 3, 2 and 4 independent experiments for BT302, BT483 and BT205 respectively. (**B**) E*x vivo* treatment schedule for res-GSC family derivation with the TMZ dose identified in (A). (**C**) E*x vivo* treatment schedule for res-GSC family derivation with a previously identified IR dose. (**D**) Stem cell frequency (fold versus the corresponding CTRL-GSC) grouped for res-GSC family member. One sample Wilcoxon test. *p= 0.0156 (**E**) Stem cell frequency in res-GSC family members, evaluated by limiting dilution assay (LDA). n = 2 independent experiments (p1), Student’s paired t test. GBM96: *p=0.043325; GBM99: *p=0.015252; GBM139: *p=0.024662. (**F**) Fisher’s exact test tables for the most relevant genetic alterations, and for MGMT promoter methylation, identified in the original GBMs. Significant association or dissociation with the ability to generate res-GSC families are indicated (red). In (A, E), data are presented as mean ± SEM.

**Fig. EV2.**
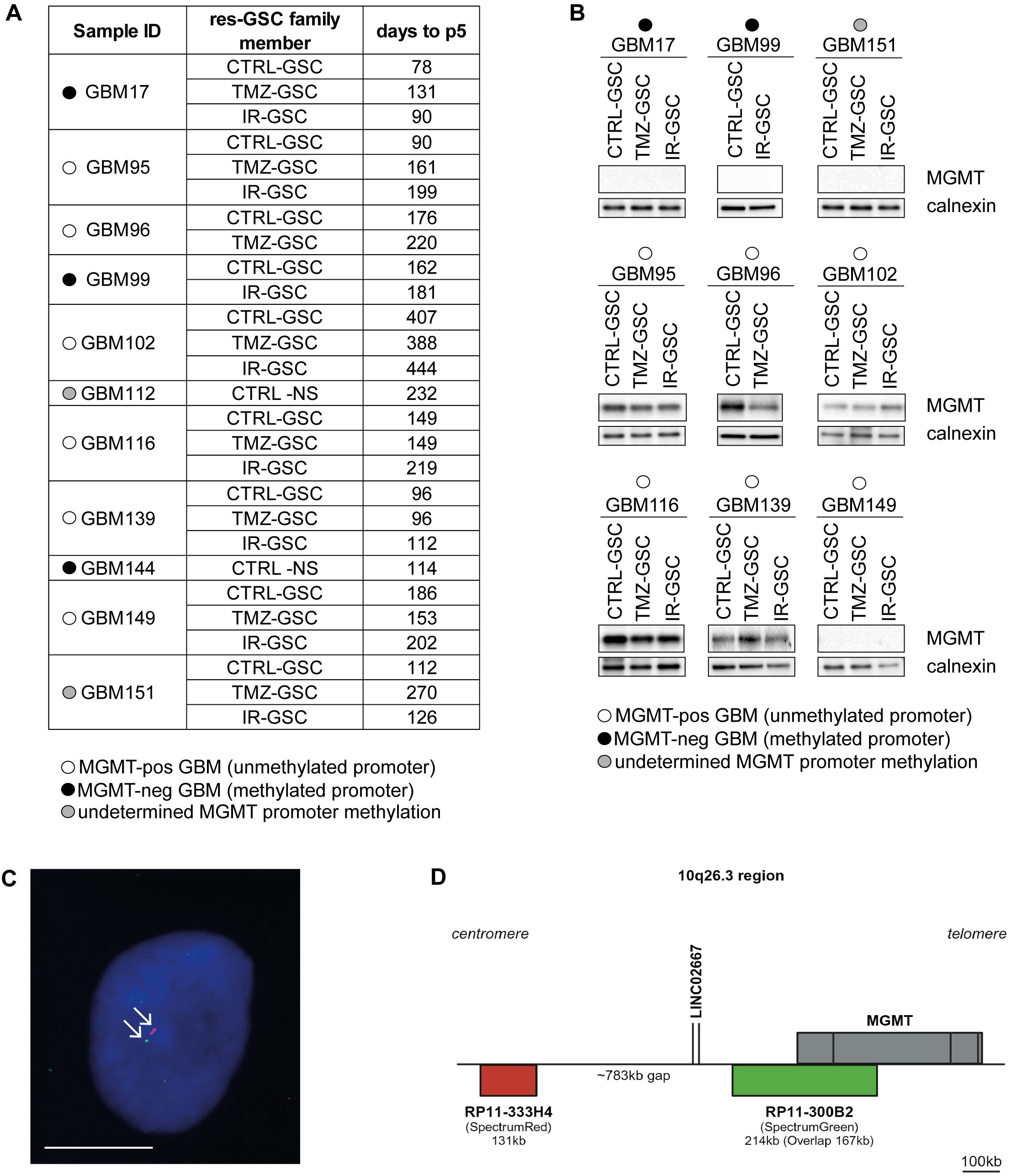
Derivation times and MGMT status of GSC family members. (**A**) Derivation times (days from GBM processing to p5) of established members in each res-GSC family. Dots indicate the MGMT promoter status of the original GBM, as detailed in Table EV3. Derivation time of TMZ-GSCs (either all cases or MGMT-pos only) vs. paired CTRL-GSCs is not statistically significantly different (Wilcoxon matched-pairs signed rank test). Derivation time of IR-GSCs vs. paired CTRL-GSCs is statistically significantly longer (Wilcoxon matched-pairs signed rank test. p=0.0078). (**B**) MGMT protein expression in all members of res-GSC families, as evaluated by western blot. Calnexin: loading control. Dots indicate the MGMT promoter status as in (A). (**C-D**) Evaluation of MGMT chromosomal localization in GBM149 CTRL-GSC evaluated by FISH. The proximity of red (RPCI11-333H4: chromosome 10 marker) and green (RPCI11-300B2: MGMT) signals allowed to exclude MGMT translocations. Scale bar: 25 µm.

**Fig. EV3.**
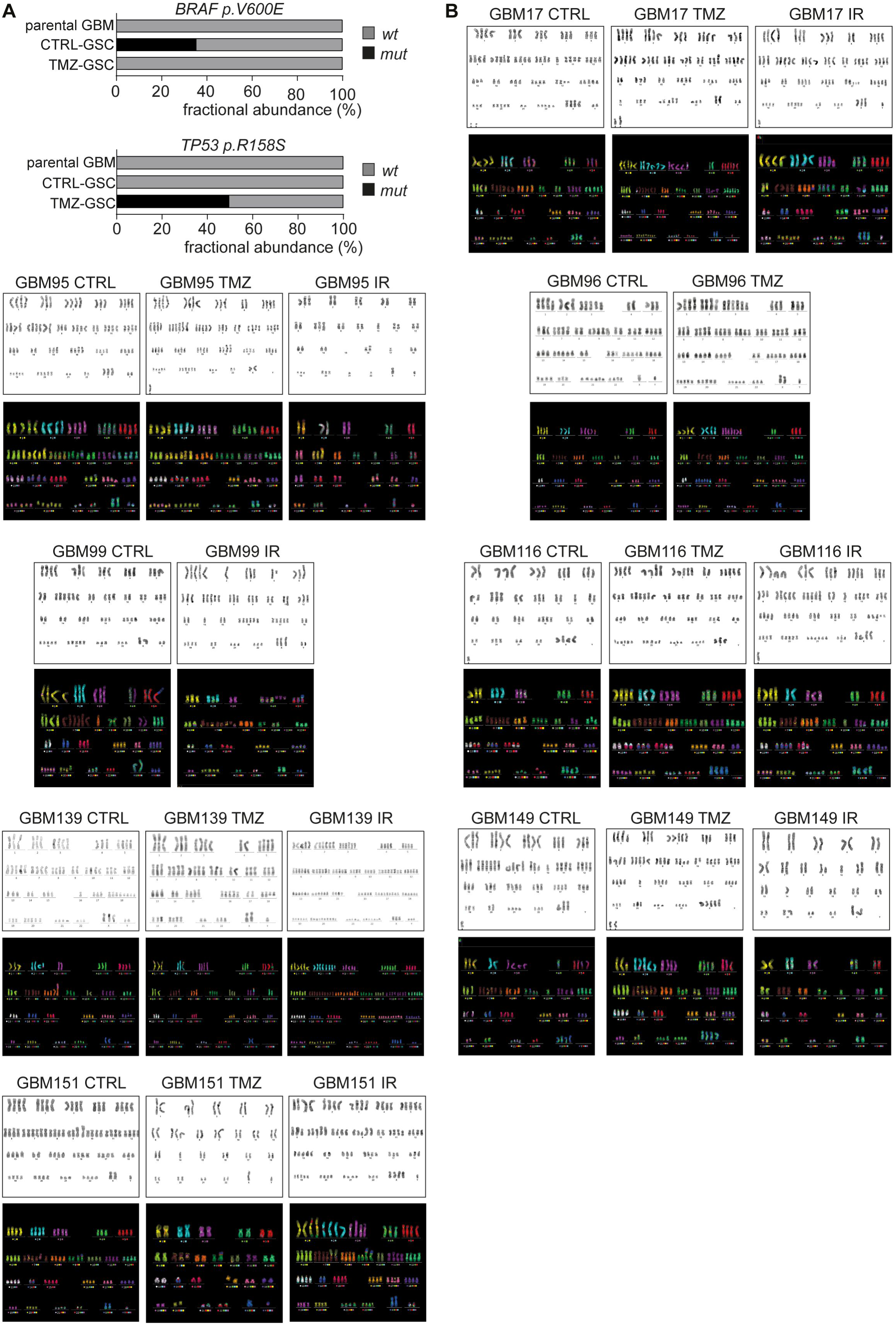
Genetics of GSCs emerging from treatments. (**A**) *BRAF* (top panel) and *TP53* (bottom panel) mutations evaluated by ddPCR in GBM149 parental GBM, CTRL-GSC and TMZ-GSC. Data are shown as fractional abundance of the indicated mutant (black) and wild type (grey) alleles. (**B**) Representative metaphase images showing chromosome number and aberrations in all res-GSC families evaluated by G-banding and M-FISH.

**Fig. EV4.**
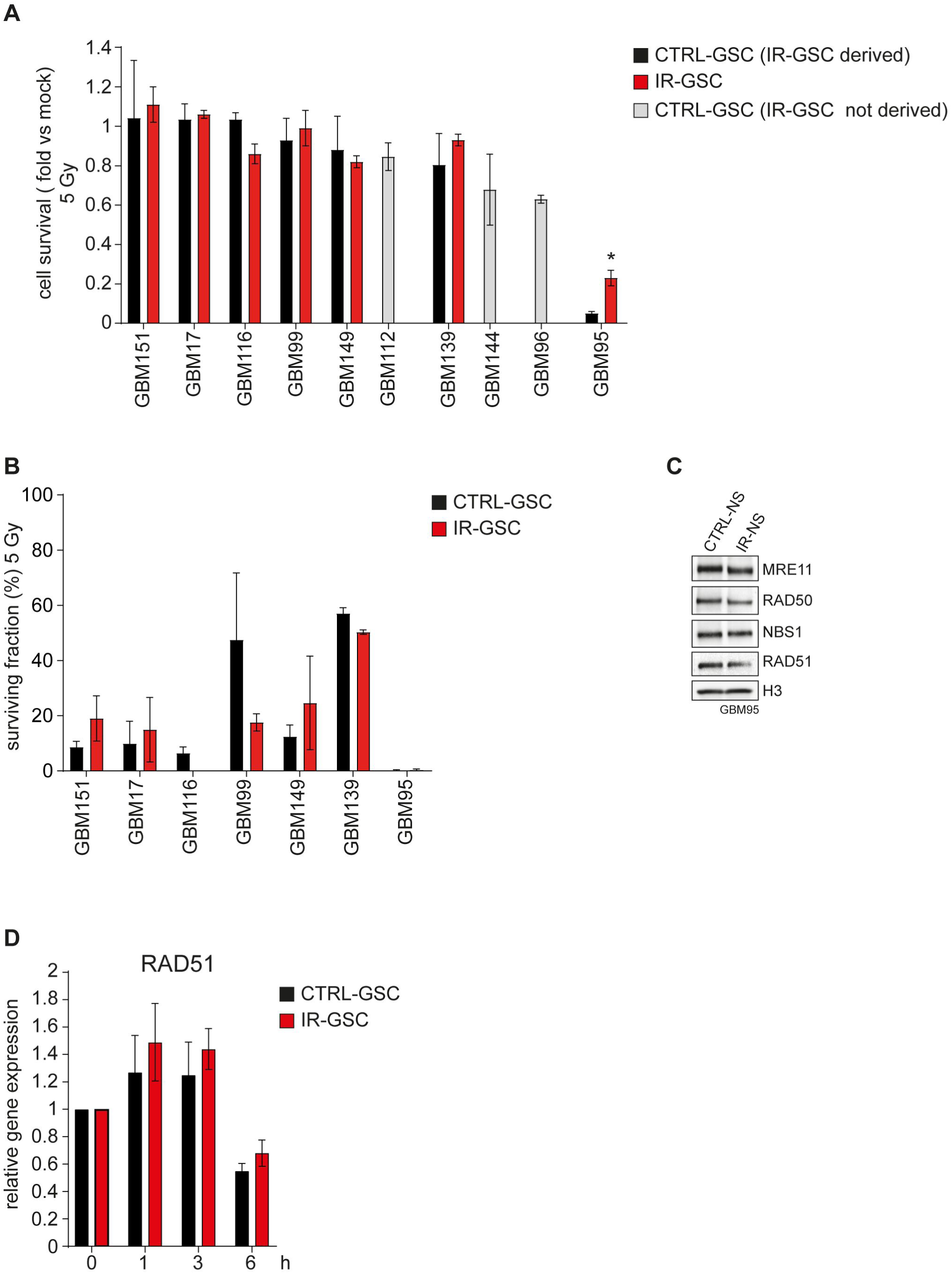
Evolution of radioresistance in GSCs under ionizing radiation pressure. (**A**) Cell survival (fold vs. mock) of CTRL-GSCs and IR-GSCs measured 48 h after irradiation with a single 5 Gy dose. n = 2 independent experiments for all cases but GBM99 (n = 3 independent experiments), Student’s paired t test. GBM95: *p= 0.041059. (**B**) Surviving fraction (%) of res-GSC families CTRL-GSC and related IR-GSC members, measured by radiobiological clonogenic assays after irradiation with a single 5 Gy dose. n = 2 independent experiments, Student’s paired t test. (**C**) Expression of the indicated proteins of the DNA damage sensor MRN complex (MRE11, RAD50 and NBS1), and RAD51 in GBM95 CTRL-GSC and IR-GSC at basal conditions. H3: loading control. (**D**) Evaluation of RAD51 gene expression in GBM95 CTRL-GSC and IR-GSC as measured by RT-PCR and shown as relative expression at the indicated time points after a single 5 Gy dose. n = 3 independent experiments, Student’s paired t test. In (A-B, D), data are presented as mean ± SEM.

**Fig. EV5.**
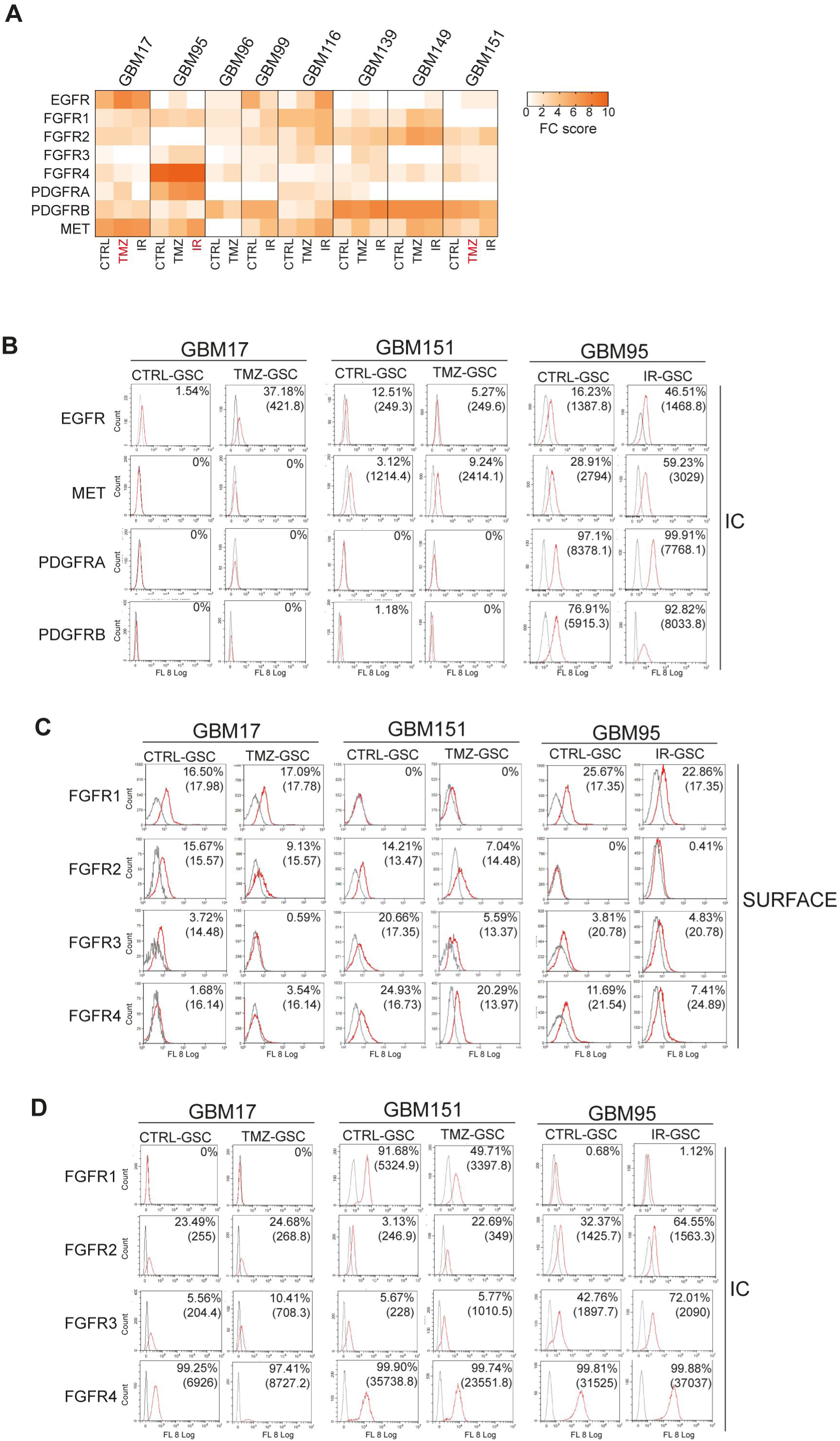
GSCs emerging from treatments exhibit changes in RTK expression and activity. (**A**) Heatmap showing flow cytometric (FC) scores of the indicated RTK surface expression (resulting from combined quantification of the percentage of positive cells and median fluorescence intensity, see Methods), in all GSCs. n = 2 independent experiments. Secondary resistant GSCs are indicated in red. (**B-D**) Flow cytometry analysis showing expression of the indicated RTKs in GBM17, GBM151 and GBM95 secondary resistant GSCs and matched CTRL-GSCs. Percentage of positive cells and mean fluorescence intensity are shown. B, D: intracytoplasmic RTK expression; C: Surface FGFR1-4 expression.

## REFERENCES

1. (2024) ISCN 2024 - An International System for Human Cytogenomic Nomenclature (2024). Cytogenet Genome Res 164: 1–224

2. Ahmed SU, Carruthers R, Gilmour L, Yildirim S, Watts C, Chalmers AJ (2015) Selective Inhibition of Parallel DNA Damage Response Pathways Optimizes Radiosensitization of Glioblastoma Stem-like Cells. Cancer Res 75: 4416–4428

3. Bao S, Wu Q, McLendon RE, Hao Y, Shi Q, Hjelmeland AB, Dewhirst MW, Bigner DD, Rich JN (2006) Glioma stem cells promote radioresistance by preferential activation of the DNA damage response. Nature 444: 756–760

4. Barthel FP, Johnson KC, Varn FS, Moskalik AD, Tanner G, Kocakavuk E, Anderson KJ, Abiola O, Aldape K, Alfaro KD et al (2019) Longitudinal molecular trajectories of diffuse glioma in adults. Nature 576: 112–120

5. Battuello P, Corti G, Bartolini A, Lorenzato A, Sogari A, Russo M, Di Nicolantonio F, Bardelli A, Crisafulli G (2024) Mutational signatures of colorectal cancers according to distinct computational workflows. Brief Bioinform 25

6. Beccuti M, Cordero F, Arigoni M, Panero R, Amparore EG, Donatelli S, Calogero RA (2018) SeqBox: RNAseq/ChIPseq reproducible analysis on a consumer game computer. Bioinformatics 34: 871–872

7. Brennan CW, Verhaak RG, McKenna A, Campos B, Noushmehr H, Salama SR, Zheng S, Chakravarty D, Sanborn JZ, Berman SH et al (2013) The somatic genomic landscape of glioblastoma. Cell 155: 462–477

8. Carro MS, Lim WK, Alvarez MJ, Bollo RJ, Zhao X, Snyder EY, Sulman EP, Anne SL, Doetsch F, Colman H et al (2010) The transcriptional network for mesenchymal transformation of brain tumours. Nature 463: 318–325

9. Cerami E, Gao J, Dogrusoz U, Gross BE, Sumer SO, Aksoy BA, Jacobsen A, Byrne CJ, Heuer ML, Larsson E et al (2012) The cBio cancer genomics portal: an open platform for exploring multidimensional cancer genomics data. Cancer Discov 2: 401–404

10. Chen J, Li Y, Yu TS, McKay RM, Burns DK, Kernie SG, Parada LF (2012) A restricted cell population propagates glioblastoma growth after chemotherapy. Nature 488: 522–526

11. Corless BC, Chang GA, Cooper S, Syeda MM, Shao Y, Osman I, Karlin-Neumann G, Polsky D (2019) Development of Novel Mutation-Specific Droplet Digital PCR Assays Detecting TERT Promoter Mutations in Tumor and Plasma Samples. J Mol Diagn 21: 274–285

12. Corti G, Bartolini A, Crisafulli G, Novara L, Rospo G, Montone M, Negrino C, Mussolin B, Buscarino M, Isella C et al (2019) A Genomic Analysis Workflow for Colorectal Cancer Precision Oncology. Clin Colorectal Cancer 18: 91–101.e103

13. Crisafulli G, Mussolin B, Cassingena A, Montone M, Bartolini A, Barault L, Martinetti A, Morano F, Pietrantonio F, Sartore-Bianchi A et al (2019) Whole exome sequencing analysis of urine trans-renal tumour DNA in metastatic colorectal cancer patients. ESMO Open 4

14. Crisafulli G, Sartore-Bianchi A, Lazzari L, Pietrantonio F, Amatu A, Macagno M, Barault L, Cassingena A, Bartolini A, Luraghi P et al (2022) Temozolomide Treatment Alters Mismatch Repair and Boosts Mutational Burden in Tumor and Blood of Colorectal Cancer Patients. Cancer Discov 12: 1656–1675

15. Darzynkiewicz Z, Crissman H, Jacobberger JW (2004) Cytometry of the cell cycle: cycling through history. Cytometry A 58: 21–32

16. De Bacco F, Casanova E, Medico E, Pellegatta S, Orzan F, Albano R, Luraghi P, Reato G, D’Ambrosio A, Porrati P et al (2012) The MET Oncogene Is a Functional Marker of a Glioblastoma Stem Cell Subtype. Cancer Res 72: 4537–4550

17. De Bacco F, D’Ambrosio A, Casanova E, Orzan F, Neggia R, Albano R, Verginelli F, Cominelli M, Poliani PL, Luraghi P et al (2016) MET inhibition overcomes radiation resistance of glioblastoma stem-like cells. EMBO Mol Med 8: 550–568

18. De Bacco F, Luraghi P, Medico E, Reato G, Girolami F, Perera T, Gabriele P, Comoglio PM, Boccaccio C (2011) Induction of MET by ionizing radiation and its role in radioresistance and invasive growth of cancer. J NatlCancer Inst 103: 645–661

19. De Bacco F, Orzan F, Casanova E, Prelli M, Boccaccio C (2023a) Protocol for in vitro establishment of heterogeneous stem-like cultures derived from whole human glioblastoma tumors. STAR Protoc 4: 102705

20. De Bacco F, Orzan F, Crisafulli G, Prelli M, Isella C, Casanova E, Albano R, Reato G, Erriquez J, D’Ambrosio A et al (2023b) Coexisting cancer stem cells with heterogeneous gene amplifications, transcriptional profiles, and malignancy are isolated from single glioblastomas. Cell Rep 42: 112816

21. Dunn J, Baborie A, Alam F, Joyce K, Moxham M, Sibson R, Crooks D, Husband D, Shenoy A, Brodbelt A et al (2009) Extent of MGMT promoter methylation correlates with outcome in glioblastomas given temozolomide and radiotherapy. Br J Cancer 101: 124–131

22. Franken NA, Rodermond HM, Stap J, Haveman J, van Bree C (2006) Clonogenic assay of cells in vitro. Nat Protoc 1: 2315–2319

23. Fu D, Calvo JA, Samson LD (2012) Balancing repair and tolerance of DNA damage caused by alkylating agents. Nat Rev Cancer 12: 104–120

24. Garofano L, Migliozzi S, Oh YT, D’Angelo F, Najac RD, Ko A, Frangaj B, Caruso FP, Yu K, Yuan J et al (2021) Pathway-based classification of glioblastoma uncovers a mitochondrial subtype with therapeutic vulnerabilities. Nat Cancer 2: 141–156

25. Gimple RC, Yang K, Halbert ME, Agnihotri S, Rich JN (2022) Brain cancer stem cells: resilience through adaptive plasticity and hierarchical heterogeneity. Nat Rev Cancer 22: 497–514

26. Hegi ME, Diserens AC, Gorlia T, Hamou MF, de TN, Weller M, Kros JM, Hainfellner JA, Mason W, Mariani L et al (2005) MGMT gene silencing and benefit from temozolomide in glioblastoma. NEnglJ Med 352: 997–1003

27. Hegi ME, Oppong FB, Perry JR, Wick W, Henriksson R, Laperriere NJ, Gorlia T, Malmström A, Weller M (2024) No benefit from TMZ treatment in glioblastoma with truly unmethylated MGMT promoter: Reanalysis of the CE.6 and the pooled Nordic/NOA-08 trials in elderly glioblastoma patients. Neuro Oncol 26: 1867–1875

28. Helderman NC, Strobel F, Bohaumilitzky L, Terlouw D, van der Werf-’t Lam AS, van Wezel T, Morreau H, von Knebel Doeberitz M, Nielsen M, Kloor M et al (2025) Lower Degree of Microsatellite Instability in Colorectal Carcinomas From MSH6-Associated Lynch Syndrome Patients. Mod Pathol 38: 100757

29. Hickman MJ, Samson LD (2004) Apoptotic signaling in response to a single type of DNA lesion, O(6)-methylguanine. Mol Cell 14: 105–116

30. Hoogstrate Y, Draaisma K, Ghisai SA, van Hijfte L, Barin N, de Heer I, Coppieters W, van den Bosch TPP, Bolleboom A, Gao Z et al (2023) Transcriptome analysis reveals tumor microenvironment changes in glioblastoma. Cancer Cell

31. Hu Y, Smyth GK (2009) ELDA: extreme limiting dilution analysis for comparing depleted and enriched populations in stem cell and other assays. J ImmunolMethods 347: 70–78

32. Johnson BE, Mazor T, Hong C, Barnes M, Aihara K, McLean CY, Fouse SD, Yamamoto S, Ueda H, Tatsuno K et al (2014) Mutational analysis reveals the origin and therapy-driven evolution of recurrent glioma. Science 343: 189–193

33. Joo KM, Jin J, Kim E, Ho KK, Kim Y, Gu KB, Kang YJ, Lathia JD, Cheong KH, Song PH et al (2012) MET signaling regulates glioblastoma stem cells. Cancer Res 72: 3828–3838

34. Juan G, Traganos F, James WM, Ray JM, Roberge M, Sauve DM, Anderson H, Darzynkiewicz Z (1998) Histone H3 phosphorylation and expression of cyclins A and B1 measured in individual cells during their progression through G2 and mitosis. Cytometry 32: 71–77

35. Kets CM, van Krieken JH, Hebeda KM, Wezenberg SJ, Goossens M, Brunner HG, Ligtenberg MJ, Hoogerbrugge N (2006) Very low prevalence of germline MSH6 mutations in hereditary non-polyposis colorectal cancer suspected patients with colorectal cancer without microsatellite instability. Br J Cancer 95: 1678–1682

36. Killela PJ, Reitman ZJ, Jiao Y, Bettegowda C, Agrawal N, Diaz LA, Friedman AH, Friedman H, Gallia GL, Giovanella BC et al (2013) TERT promoter mutations occur frequently in gliomas and a subset of tumors derived from cells with low rates of self-renewal. Proc Natl Acad Sci U S A 110: 6021–6026

37. Kim H, Zheng S, Amini SS, Virk SM, Mikkelsen T, Brat DJ, Grimsby J, Sougnez C, Muller F, Hu J et al (2015) Whole-genome and multisector exome sequencing of primary and post-treatment glioblastoma reveals patterns of tumor evolution. Genome Res 25: 316–327

38. Kim KH, Migliozzi S, Koo H, Hong JH, Park SM, Kim S, Kwon HJ, Ha S, Garofano L, Oh YT et al (2024) Integrated proteogenomic characterization of glioblastoma evolution. Cancer Cell 42: 358–377.e358

39. Kim Y, Varn FS, Park SH, Yoon BW, Park HR, Lee C, Verhaak RGW, Paek SH (2021) Perspective of mesenchymal transformation in glioblastoma. Acta Neuropathol Commun 9: 50

40. Kocakavuk E, Anderson KJ, Varn FS, Johnson KC, Amin SB, Sulman EP, Lolkema MP, Barthel FP, Verhaak RGW (2021) Radiotherapy is associated with a deletion signature that contributes to poor outcomes in patients with cancer. Nat Genet 53: 1088–1096

41. Li H, Wu Y, Chen Y, Lv J, Qu C, Mei T, Zheng Y, Ye C, Li F, Ge S et al (2025) Overcoming temozolomide resistance in glioma: recent advances and mechanistic insights. Acta Neuropathol Commun 13: 126

42. Liau BB, Sievers C, Donohue LK, Gillespie SM, Flavahan WA, Miller TE, Venteicher AS, Hebert CH, Carey CD, Rodig SJ et al (2017) Adaptive Chromatin Remodeling Drives Glioblastoma Stem Cell Plasticity and Drug Tolerance. Cell Stem Cell 20: 233–246.e237

43. Liu J, Cao S, Imbach KJ, Gritsenko MA, Lih TM, Kyle JE, Yaron-Barir TM, Binder ZA, Li Y, Strunilin I et al (2024) Multi-scale signaling and tumor evolution in high-grade gliomas. Cancer Cell 42: 1217–1238.e1219

44. Liu R, Chen Y, Liu G, Li C, Song Y, Cao Z, Li W, Hu J, Lu C, Liu Y (2020) PI3K/AKT pathway as a key link modulates the multidrug resistance of cancers. Cell Death Dis 11: 797

45. Louis DN, Perry A, Wesseling P, Brat DJ, Cree IA, Figarella-Branger D, Hawkins C, Ng HK, Pfister SM, Reifenberger G et al (2021) The 2021 WHO Classification of Tumors of the Central Nervous System: a summary. Neuro Oncol 23: 1231–1251

46. Lucas CG, Al-Adli NN, Young JS, Gupta R, Morshed RA, Wu J, Ravindranathan A, Shai A, Oberheim Bush NA, Taylor JW et al (2025) Longitudinal multimodal profiling of IDH-wildtype glioblastoma reveals the molecular evolution and cellular phenotypes underlying prognostically different treatment responses. Neuro Oncol 27: 89–105

47. Lucchini S, Nicholson JG, Zhang X, Househam J, Lim YM, Mossner M, Millner TO, Brandner S, Graham T, Marino S (2025) A novel model of glioblastoma recurrence to identify therapeutic vulnerabilities. EMBO Mol Med 17: 1325–1354

48. Meyer M, Reimand J, Lan X, Head R, Zhu X, Kushida M, Bayani J, Pressey JC, Lionel AC, Clarke ID et al (2015) Single cell-derived clonal analysis of human glioblastoma links functional and genomic heterogeneity. Proc Natl Acad Sci U S A 112: 851–856

49. Morales-Juarez DA, Jackson SP (2022) Clinical prospects of WRN inhibition as a treatment for MSI tumours. NPJ Precis Oncol 6: 85

50. Nagel ZD, Margulies CM, Chaim IA, McRee SK, Mazzucato P, Ahmad A, Abo RP, Butty VL, Forget AL, Samson LD (2014) Multiplexed DNA repair assays for multiple lesions and multiple doses via transcription inhibition and transcriptional mutagenesis. Proc Natl Acad Sci U S A 111: E1823–1832

51. Neftel C, Laffy J, Filbin MG, Hara T, Shore ME, Rahme GJ, Richman AR, Silverbush D, Shaw ML, Hebert CM et al (2019) An Integrative Model of Cellular States, Plasticity, and Genetics for Glioblastoma. Cell 178: 835–849.e821

52. Nicholson JG, Fine HA (2021) Diffuse Glioma Heterogeneity and Its Therapeutic Implications. Cancer Discov 11: 575–590

53. Oldrini B, Vaquero-Siguero N, Mu Q, Kroon P, Zhang Y, Galán-Ganga M, Bao Z, Wang Z, Liu H, Sa JK et al (2020) MGMT genomic rearrangements contribute to chemotherapy resistance in gliomas. Nat Commun 11: 3883

54. Orzan F, De Bacco F, Crisafulli G, Pellegatta S, Mussolin B, Siravegna G, D’Ambrosio A, Comoglio PM, Finocchiaro G, Boccaccio C (2017) Genetic Evolution of Glioblastoma Stem-Like Cells From Primary to Recurrent Tumor. Stem Cells 35: 2218–2228

55. Ostermann S, Csajka C, Buclin T, Leyvraz S, Lejeune F, Decosterd LA, Stupp R (2004) Plasma and cerebrospinal fluid population pharmacokinetics of temozolomide in malignant glioma patients. Clin Cancer Res 10: 3728–3736

56. Osuka S, Van Meir EG (2017) Overcoming therapeutic resistance in glioblastoma: the way forward. J Clin Invest 127: 415–426

57. Ozawa T, Brennan CW, Wang L, Squatrito M, Sasayama T, Nakada M, Huse JT, Pedraza A, Utsuki S, Yasui Y et al (2010) PDGFRA gene rearrangements are frequent genetic events in PDGFRA-amplified glioblastomas. Genes Dev 24: 2205–2218

58. Pagani F, Orzan F, Lago S, De Bacco F, Prelli M, Cominelli M, Somenza E, Gryzik M, Balzarini P, Ceresa D et al (2025) Concurrent RB1 and P53 pathway disruption predisposes to the development of a primitive neuronal component in high-grade gliomas depending on MYC-driven EBF3 transcription. Acta Neuropathol 149: 8

59. Piccirillo SG, Colman S, Potter NE, van Delft FW, Lillis S, Carnicer MJ, Kearney L, Watts C, Greaves M (2015) Genetic and functional diversity of propagating cells in glioblastoma. Stem Cell Reports 4: 7–15

60. Piccirillo SG, Combi R, Cajola L, Patrizi A, Redaelli S, Bentivegna A, Baronchelli S, Maira G, Pollo B, Mangiola A et al (2009) Distinct pools of cancer stem-like cells coexist within human glioblastomas and display different tumorigenicity and independent genomic evolution. Oncogene 28: 1807–1811

61. Piett CG, Pecen TJ, Laverty DJ, Nagel ZD (2021) Large-scale preparation of fluorescence multiplex host cell reactivation (FM-HCR) reporters. Nat Protoc 16: 4265–4298

62. Pignochino Y, Crisafulli G, Giordano G, Merlini A, Berrino E, Centomo ML, Chiabotto G, Brusco S, Basiricò M, Maldi E et al (2021) PARP1 Inhibitor and Trabectedin Combination Does Not Increase Tumor Mutational Burden in Advanced Sarcomas-A Preclinical and Translational Study. Cancers (Basel) 13

63. Prat M, Crepaldi T, Pennacchietti S, Bussolino F, Comoglio PM (1998) Agonistic monoclonal antibodies against the Met receptor dissect the biological responses to HGF. J Cell Sci 111 (Pt 2): 237–247

64. Prigent C, Dimitrov S (2003) Phosphorylation of serine 10 in histone H3, what for? J Cell Sci 116: 3677–3685

65. Pu Y, Li L, Peng H, Liu L, Heymann D, Robert C, Vallette F, Shen S (2023) Drug-tolerant persister cells in cancer: the cutting edges and future directions. Nat Rev Clin Oncol 20: 799–813

66. Rondon MR, Raffel SJ, Goodman RM, Handelsman J (1999) Toward functional genomics in bacteria: analysis of gene expression in Escherichia coli from a bacterial artificial chromosome library of Bacillus cereus. Proc Natl Acad Sci U S A 96: 6451–6455

67. Russo M, Chen M, Mariella E, Peng H, Rehman SK, Sancho E, Sogari A, Toh TS, Balaban NQ, Batlle E et al (2024) Cancer drug-tolerant persister cells: from biological questions to clinical opportunities. Nat Rev Cancer 24: 694–717

68. Salem ME, Bodor JN, Puccini A, Xiu J, Goldberg RM, Grothey A, Korn WM, Shields AF, Worrilow WM, Kim ES et al (2020) Relationship between MLH1, PMS2, MSH2 and MSH6 gene-specific alterations and tumor mutational burden in 1057 microsatellite instability-high solid tumors. Int J Cancer 147: 2948–2956

69. Schulte A, Gunther HS, Martens T, Zapf S, Riethdorf S, Wulfing C, Stoupiec M, Westphal M, Lamszus K (2012) Glioblastoma stem-like cell lines with either maintenance or loss of high-level EGFR amplification, generated via modulation of ligand concentration. Clin Cancer Res 18: 1901–1913

70. Schulze Heuling E, Knab F, Radke J, Eskilsson E, Martinez-Ledesma E, Koch A, Czabanka M, Dieterich C, Verhaak RG, Harms C et al (2017) Prognostic Relevance of Tumor Purity and Interaction with MGMT Methylation in Glioblastoma. Mol Cancer Res 15: 532–540

71. Shaw RJ, Liloglou T, Rogers SN, Brown JS, Vaughan ED, Lowe D, Field JK, Risk JM (2006) Promoter methylation of P16, RARbeta, E-cadherin, cyclin A1 and cytoglobin in oral cancer: quantitative evaluation using pyrosequencing. Br J Cancer 94: 561–568

72. Sjöstedt E, Zhong W, Fagerberg L, Karlsson M, Mitsios N, Adori C, Oksvold P, Edfors F, Limiszewska A, Hikmet F et al (2020) An atlas of the protein-coding genes in the human, pig, and mouse brain. Science 367

73. Sloan AR, Silver DJ, Kint S, Gallo M, Lathia JD (2024) Cancer stem cell hypothesis 2.0 in glioblastoma: Where are we now and where are we going? Neuro Oncol 26: 785–795

74. Sondka Z, Dhir NB, Carvalho-Silva D, Jupe S, Madhumita, McLaren K, Starkey M, Ward S, Wilding J, Ahmed M et al (2024) COSMIC: a curated database of somatic variants and clinical data for cancer. Nucleic Acids Res 52: D1210–D1217

75. Spitzer A, Johnson KC, Nomura M, Garofano L, Nehar-Belaid D, Darnell NG, Greenwald AC, Bussema L, Oh YT, Varn FS et al (2025) Deciphering the longitudinal trajectories of glioblastoma ecosystems by integrative single-cell genomics. Nat Genet 57: 1168–1178

76. Squatrito M, Holland EC (2011) DNA damage response and growth factor signaling pathways in gliomagenesis and therapeutic resistance. Cancer Res 71: 5945–5949

77. Steinhaus R, Proft S, Schuelke M, Cooper DN, Schwarz JM, Seelow D (2021) MutationTaster2021. Nucleic Acids Res 49: W446–W451

78. Stupp R, Mason WP, van den Bent MJ, Weller M, Fisher B, Taphoorn MJ, Belanger K, Brandes AA, Marosi C, Bogdahn U et al (2005) Radiotherapy plus concomitant and adjuvant temozolomide for glioblastoma. N Engl J Med 352: 987–996

79. Suvà ML, Tirosh I (2020) The Glioma Stem Cell Model in the Era of Single-Cell Genomics. Cancer Cell 37: 630–636

80. Szerlip NJ, Pedraza A, Chakravarty D, Azim M, McGuire J, Fang Y, Ozawa T, Holland EC, Huse JT, Jhanwar S et al (2012) Intratumoral heterogeneity of receptor tyrosine kinases EGFR and PDGFRA amplification in glioblastoma defines subpopulations with distinct growth factor response. Proc Natl Acad Sci U S A 109: 3041–3046

81. Tanner G, Barrow R, Ajaib S, Al-Jabri M, Ahmed N, Pollock S, Finetti M, Rippaus N, Bruns AF, Syed K et al (2024) IDHwt glioblastomas can be stratified by their transcriptional response to standard treatment, with implications for targeted therapy. Genome Biol 25: 45

82. Tordai H, Torres O, Csepi M, Padányi R, Lukács GL, Hegedűs T (2024) Analysis of AlphaMissense data in different protein groups and structural context. Sci Data 11: 495

83. Touat M, Li YY, Boynton AN, Spurr LF, Iorgulescu JB, Bohrson CL, Cortes-Ciriano I, Birzu C, Geduldig JE, Pelton K et al (2020) Mechanisms and therapeutic implications of hypermutation in gliomas. Nature 580: 517–523

84. Varn FS, Johnson KC, Martinek J, Huse JT, Nasrallah MP, Wesseling P, Cooper LAD, Malta TM, Wade TE, Sabedot TS et al (2022) Glioma progression is shaped by genetic evolution and microenvironment interactions. Cell 185: 2184–2199.e2116

85. Verhaak RG, Hoadley KA, Purdom E, Wang V, Qi Y, Wilkerson MD, Miller CR, Ding L, Golub T, Mesirov JP et al (2010) Integrated genomic analysis identifies clinically relevant subtypes of glioblastoma characterized by abnormalities in PDGFRA, IDH1, EGFR, and NF1. Cancer Cell 17: 98–110

86. Wang J, Cazzato E, Ladewig E, Frattini V, Rosenbloom DI, Zairis S, Abate F, Liu Z, Elliott O, Shin YJ et al (2016) Clonal evolution of glioblastoma under therapy. Nat Genet 48: 768–776

87. Wang L, Jung J, Babikir H, Shamardani K, Jain S, Feng X, Gupta N, Rosi S, Chang S, Raleigh D et al (2022) A single-cell atlas of glioblastoma evolution under therapy reveals cell-intrinsic and cell-extrinsic therapeutic targets. Nat Cancer 3: 1534–1552

88. Wang Q, Hu B, Hu X, Kim H, Squatrito M, Scarpace L, deCarvalho AC, Lyu S, Li P, Li Y et al (2017) Tumor Evolution of Glioma-Intrinsic Gene Expression Subtypes Associates with Immunological Changes in the Microenvironment. Cancer Cell 32: 42–56.e46

89. Weller M, Wen PY, Chang SM, Dirven L, Lim M, Monje M, Reifenberger G (2024) Glioma. Nat Rev Dis Primers 10: 33

90. Wen PY, Weller M, Lee EQ, Touat M, Khasraw M, Rahman R, Platten M, Lim M, Winkler F, Horbinski C et al (2025) Glioblastoma in Adults: A Society for Neuro-Oncology (SNO) and European Society of Neuro-Oncology (EANO) Consensus Review on Current Management and Future Directions. Neuro Oncol

91. Woo PYM, Li Y, Chan AHY, Ng SCP, Loong HHF, Chan DTM, Wong GKC, Poon W-S, 2019. A multifaceted review of temozolomide resistance mechanisms in glioblastoma beyond O-6-methylguanine-DNA methyltransferase. Glioma.

92. Zhang J, Feng Y, Li G, Zhang X, Zhang Y, Qin Z, Zhuang D, Qiu T, Shi Z, Zhu W et al (2023) Distinct aneuploid evolution of astrocytoma and glioblastoma during recurrence. NPJ Precis Oncol 7: 97

